# Single-molecule visualisation of human Hsp70-driven conformational remodelling during stress

**DOI:** 10.64898/2026.06.01.729260

**Authors:** Bailey Skewes, Shannon McMahon, Nicola K. Auld, Nicholas R. Marzano, Antoine M. van Oijen, Heath Ecroyd

## Abstract

The Hsp70 chaperone system plays a central role in the folding of nascent polypeptides and in preventing protein misfolding and aggregation during cellular stress. However, the precise mechanism by which the human Hsp70, HspA8, remodels the conformations of individual chemically misfolded clients remains unclear. Moreover, it is unknown whether this mechanism differs from that used by Hsp70 to engage clients during heat stress to preserve native function. To address these questions, we report here the use of single-molecule fluorescence resonance energy transfer (smFRET) to temporally interrogate how the human Hsp70 system regulates the conformation of a heat-sensitive client protein, firefly luciferase (Fluc), following chemical denaturation and during heat stress. We find that Hsp70 recognises both chemically denatured and heat-induced misfolded states of Fluc and resolves them by conformational expansion. Release from a Hsp70-bound state, a process driven by the nucleotide exchange factor, Hsp110, guides Fluc toward productive folding trajectories that would otherwise be unlikely to occur spontaneously following collapse from a conformationally unfolded state. Moreover, we demonstrate that both temperature and the conformational state of misfolded Fluc dictate the ability of HspA8 to meaningfully resolve non-native structure within the protein. Collectively, this work provides direct visualisation of the mechanisms by which Hsp70 modulates client conformations under diverse stress conditions to preserve proteome integrity.

## Introduction

Hsp70 serves as a central hub within the molecular chaperone network, coordinating a myriad of roles within the cell that include co-translational folding, refolding of misfolded proteins and aggregate disassembly, processes that are essential for the maintenance of cellular protein homeostasis (proteostasis)^1^. Hsp70 is highly conserved across all orders of life and comprises of an N-terminal nucleotide binding domain (NBD), which binds and hydrolyses ATP, allosterically linked via a short linker to the client binding domain (CBD). The CBD recognises short hydrophobic 7-8 amino acid motifs that typically become solvent exposed in misfolded protein clients^2^. Hsp70 displays a nucleotide-dependent affinity for client proteins, a process tightly regulated by its co-chaperones Hsp40 and nucleotide exchange factors (NEFs). In the canonical model of Hsp70 function, Hsp40 first recognises misfolded client proteins and delivers them to Hsp70 in its ATP-bound state, whereby the CBD resides in an open conformation characterised by rapid rates of client binding and release. ATP hydrolysis is accelerated by Hsp40, which allosterically triggers a conformational change in Hsp70 to the ‘closed’ ADP-bound state in which the client is stably captured within the CBD^3^. Finally, association of a NEF with the Hsp70-client complex promotes exchange of ADP for ATP in solution, which returns Hsp70 to the low affinity ‘open’ state, facilitating client release and providing an opportunity for the client to fold into the native state.

Studies of the archetypal *Escherichia coli* (*E. coli*) Hsp70 system have demonstrated that binding of Hsp70 (DnaK in *E. coli*) to client proteins imparts an unfolding force such that they enter expanded conformational states^4–7^. One proposed explanation for the origin of this force is ‘entropic pulling’, whereby the stable binding of multiple bulky Hsp70 molecules restricts the conformational freedom of the client-chaperone complex through excluded volume effects. This energetic constraint can be resolved by Hsp70 ‘pulling’ the client into a more extended conformation^8–10^. However, it remains unresolved whether Hsp70 actively contributes to the subsequent folding of the client following release, or whether it primarily functions as an ‘unfoldase’. Moreover, it is unclear whether the human Hsp70 chaperone machinery (which has evolved a diverse array of Hsp70s, Hsp40s and NEFs^11,12^) promotes protein refolding via the same underlying mechanism as observed for the single isoforms present in *E. coli*.

Most *in vitro* studies of Hsp70-mediated refolding have focused on the capacity of the chaperone system to refold misfolded client proteins generated by dilution from chemical denaturant^4–7^. However, such denaturing conditions are not representative of the types of stressors that typically threaten proteostasis in the cell. Notably, misfolded states generated by chemical denaturation often differ substantially from those produced by more physiologically relevant stressors such as heat, which commonly induce localised and partial unfolding rather than the global loss of structure characteristic of chemical denaturation^13,14^. In addition, these experiments are often performed post-denaturation, once the protein population has already become completely misfolded; a scenario that does not reflect cellular conditions whereby folding and misfolding generate a heterogenous ensemble of conformations. As such, studies following chemical denaturation of the client only allows the interrogation of how the Hsp70 system *refolds* globally misfolded client proteins rather than addressing the ability of the chaperone system to *maintain* clients in their native state. Consistent with this latter function, the Hsp70 chaperone system has been found to be protective against heat denaturation of client proteins, effectively maintaining them in a biologically active state under conditions that would typically result in their misfolding and aggregation^5,15^. However, the molecular mechanisms by which the Hsp70 system maintains native states remain unclear; specifically, does Hsp70 employ the same mechanisms to maintain a native state compared to refolding of a misfolded protein? Additionally, what are the key kinetic and thermodynamic principles that underlie these processes?

Establishing the mechanisms by which Hsp70 exerts these functions is complicated by the heterogeneity of client-chaperone complexes and the presence of competing reactions, such as client aggregation, which are masked in traditional ensemble-based approaches. To overcome these limitations, we utilise single molecule fluorescence resonance energy transfer (smFRET) to temporally monitor the conformational dynamics of individual firefly luciferase (Fluc) proteins as they are refolded from distinct misfolded states by a human Hsp70 chaperone system comprising the constitutively expressed Hsp70 (HspA8), Hsp40 (DnaJA2) and NEF (Hsp110). Additionally, we established an immobilised smFRET assay under heat-destabilising conditions that enables the interrogation of Hsp70-mediated maintenance of a native protein population without competing aggregation reactions.

We find that HspA8 resolves misfolded Fluc by actively unfolding the client into an expanded conformation, with both the stability and extent of unfolding dependent on the presence of NEF. In comparison, HspA8 only transiently unfolds misfolded client proteins under heat-stress conditions, a process sufficient to maintain a predominantly native state population. Notably, these transient interactions are not sufficient to restore chemically misfolded Fluc to a native state, which instead requires prolonged periods of conformational expansion (likely correlated with increased Hsp70 binding) to effectively resolve more globally misfolded states and promote folding. Interestingly, client refolding from expanded conformational states (induced by Hsp70) during heat stress produced alternatively misfolded Fluc states not always observed in the absence of Hsp70, indicating that molecular chaperones can inadvertently direct clients down alternate misfolding pathways. Collectively, we define the HspA8-mediated refolding pathway of a chemically and heat-induced misfolded client protein and demonstrate that HspA8 interacts with these alternate misfolded conformers with distinct kinetic and thermodynamic behaviours.

## Methods Materials

Plasmids encoding for human chaperone proteins (HspA8, DnaJA2, Hsp110) were kindly donated by Dr Nadinath Nillegoda (Monash University, Australia).

### Expression and purification of recombinant protein

In order to monitor the interdomain conformation of Fluc via smFRET, a construct containing two cysteine substitutions (K141C, K491C) for site-specific labelling was generated, referred to as Fluc^IDS^ (interdomain sensor). Fluc^IDS^ was expressed and purified as previously described^6^ with the following modification. During initial purification of the cell lysate with immobilized metal affinity chromatography (IMAC), column-bound Fluc was incubated with 50 µM D-Biotin in 10 mM bicine buffer (pH 8.3) to promote dissociation of non-specifically bound, co-expressed biotin ligase.

HspA8 and Hsp110 were expressed and purified as described previously^16^. Expression of DnaJA2 was performed by selecting *E. coli* (BL21) Rosetta™ cells transformed with the plasmid encoding DnaJA2 to inoculate 100 mL starter cultures of 2×YT media (1.6% [w/v] tryptone, 1% [w/v] yeast extract, 0.5% [w/v] NaCl, pH 7.0) supplemented with kanamycin (50 µg/mL) and chloramphenicol (25 µg/mL) and grown overnight at 37 °C with constant agitation in an orbital shaker at 180 rpm. Starter cultures were added to 1 L expression cultures of 2×YT supplemented with only kanamycin (50 µg/mL) and incubated at 30 °C at 180 rpm until reaching an optical density at 600 nm of 1.0. Protein expression was induced by addition of 0.5 mM isopropyl-β-D-1-thiogalactopyranoside and incubation at 30 °C for 4 h prior to harvesting of pellets by centrifugation (5,000 × *g*, 10 min, 4 °C). Pellets were stored at-20 °C until extraction of recombinant protein.

The bacterial pellet containing expressed DnaJA2 was resuspended in ice-cold lysis buffer (50 mM HEPES-KOH [pH 7.4], 5 mM MgCl_2_, 2 mM β-mercaptoethanol [BME], 20 mM imidazole, 10% glycerol [v/v], 1 x cOmplete^TM^ EDTA-free protease inhibitor cocktail [Thermo Fisher Scientific, USA]). Lysates were then supplemented with DNase I (3.0 µg/mL), lysozyme (0.5 mg/mL) and 500 mM KCl. Following 30 min of incubation at 4 °C, the cell lysate was then probe sonicated for 3 min (10 s on/20 s off) at 45% amplitude and centrifuged twice at 24,000 × *g* for 20 min at 4 °C to remove cell debris. The clarified lysate was then passed through a 0.45 µm filter prior to purification.

The bacterial cell lysate (∼ 50 mL) was loaded onto a 5 mL HisTrap^TM^ HP Sepharose™ column (Cytiva) pre-equilibrated in lysis buffer and washed with Buffer A (50 mM HEPES-KOH [pH 7.4], 5 mM MgCl_2_, 2 mM BME, 20 mM imidazole, 10% [v/v] glycerol, 500 mM KCl) until the absorbance at 280 nm returned to baseline. Column-bound protein was eluted upon addition of elution buffer (Buffer A supplemented with 300 mM imidazole). Fractions containing the protein of interest were pooled and supplemented with 6 × His-tagged ULP1 protease (4 µg/mL per mg of recombinant protein) prior to overnight dialysis at 4 °C into cleavage buffer (50 mM HEPES-KOH [pH 7.4], 5 mM MgCl_2_, 2 mM BME, 20 mM imidazole, 100 mM KCl, 10% [v/v] glycerol). Cleaved proteins were loaded onto the HisTrap^TM^ HP Sepharose in the same manner as described above to separate recombinant DnaJA2 from the 6 × His-SUMO tag. Cleaved proteins were loaded onto a HiLoad^®^ 16/600 Superdex^®^ 200 size-exclusion column (Cytiva) pre-equilibrated in 50 mM HEPES-KOH (pH 7.4), 5 mM MgCl_2_, 2 mM BME, 10% glycerol (v/v) and 500 mM KCl. Recombinant DnaJA2 was eluted from the column at a flowrate of 1.5 mL/minute into 10 mL fractions. Fractions containing recombinant protein were pooled, concentrated using a 10 kDa MWCO Pierce^TM^ Protein Concentrator (Thermo Fisher Scientific) and dialysed overnight at 4 °C into anion exchange chromatography (AEC) buffer A (50 mM HEPES-KOH [pH 7.4], 10 mM KCl, 5 mM MgCl_2_, 1 mM BME, 10% [v/v] glycerol) prior to subsequent anion-exchange purification. Dialysate was loaded onto a 1 mL Mono Q^TM^ (Cytiva) column pre-equilibrated in AEC buffer A. Bound protein was eluted from the column via a linear salt gradient (10 – 500 mM KCl) across 20 column volumes. Protein-containing fractions were once again pooled, concentrated as above and concentration determined by use of a NanoDrop^TM^ 2000c Spectrophometer (Thermo Fisher Scientific) prior to storage at-80 °C.

### Fluc luminescence assays

To confirm the ability of molecular chaperones to refold a denatured client, luminescence assays were performed. Native Fluc^IDS^ was chemically denatured by incubation in unfolding buffer (50 mM HEPES-KOH [pH 7.5], 50 mM KCl, 5 mM MgCl_2_, 2 mM dithiothreitol [DTT], 4 M guanidium hydrochloride [GdHCl]) for 10 min at room temperature. Refolding was initiated by a 1:100 dilution of chemically-denatured Fluc^IDS^ into refolding buffer (50 mM HEPES-KOH [pH 7.5], 50 mM KCl, 5 mM MgCl_2_, 2 mM DTT, 0.05% [v/v] Tween-20, 5 mM ATP) to a final concentration of 10 nM. Throughout the refolding reaction, aliquots were taken and added to a white 96-well Costar^®^ plate (Sigma-Aldrich, USA). Luminescence was recorded for 5 s following an automated injection of a 10-fold excess of luciferin buffer (25 mM glycylglycine [pH 7.4], 0.25 mM luciferin, 100 mM KCl, 2 mM ATP) using a POLARstar^®^ Omega (BMG Labtech, Germany) plate reader. Measurements for chemically-denatured refolding reactions were performed at room temperature. For chaperone-mediated refolding reactions, chemically-denatured Fluc^IDS^ was diluted into refolding buffer supplemented with either 5 µM HspA8 and 2 µM DnaJA2 alone (A8/A2) or supplemented with 0.5 µM Hsp110 (A8/A2/110). The luminescence values were normalised to the signal from that measured for native Fluc^IDS^ and the data were fit to one-phase association curves.

To determine how molecular chaperones act to maintain a native state in the presence of denaturing conditions, refolding assays were also performed at a higher temperature (37 °C) which was previously demonstrated to cause Fluc misfolding^5,17^. For these experiments, native Fluc^IDS^ (10 nM) was incubated in refolding buffer at 37 °C in a QuantStudio^TM^ 5 Real-Time PCR System (Thermo Fisher Scientific) to ensure adequate temperature control was maintained. Changes in activity were monitored and normalised to the initial luminescence value (*i.e*., *t* = 0 h) for each replicate. The reaction was incubated for 2 h either in the absence (Fluc^IDS^ alone) or presence of 3 µM HspA8, 2 µM DnaJA2, 0.5 µM Hsp110 and 5 mM ATP (A8/A2/110). After 2 h, the reaction was returned to room temperature, and the enzymatic activity was monitored for an additional hour. For all reactions, aliquots were taken at various timepoints and luminescence measured as described above, with readings performed at the same temperature in which they had been incubated previously (*i.e*., either 37 °C or 25 °C).

### Protein labelling

Fluc^IDS^ was labelled with fluorescent dyes as described previously^18^ with minor modifications. Briefly, Fluc^IDS^ (2 mg/mL) was incubated in the presence of 5 mM tris(2-carboxyethyl)phosphine and 40% (w/v) ammonium sulfate whilst rotating for 1 h at 4 °C to reduce disulfide bonds and ensure access to cysteines for labelling. Protein was then pelleted by centrifugation at 20,000 × *g* for 5 min prior to resuspension in degassed labelling buffer (100 mM Na_2_PO_4_ [pH 7.4], 200 mM NaCl, 1 mM EDTA) supplemented with 40% (w/v) ammonium sulfate. Fluc^IDS^ was then pelleted again at 20,000 × *g* for 15 min and resuspended in degassed labelling buffer. Fluc^IDS^ was then incubated in the presence of a four-fold excess of maleimide-functionalised Alexa Fluor 555 (AF555) (Thermo Fisher Scientific) and a six-fold excess of maleimide-functionalised Alexa Fluor 647 (AF647) (Thermo Fisher Scientific) to permit dual labelling. The reaction mix was incubated on a rotator overnight at 4 °C before excess dye was removed by multiple passes over a 7,000 Da MWCO Zeba™ Spin Desalting Column (Thermo Fisher Scientific) pre-equilibrated in 50 mM Tris (pH 7.5) supplemented with 20% (v/v) glycerol. The concentration and degree of labelling was determined using the NanoDrop^TM^ 2000c Spectrophometer.

### TIRF microscope setup

Samples were imaged on a custom-built TIRF microscopy setup, employing the use of an inverted optical microscope (Nikon Eclipse TI-2) with an electron-multiplied charge-coupled device (emCCD) camera (Andor iXon Life 897, Oxford Instruments, UK). Diode-pumped, continuous-wave solid-state lasers (200 mW Sapphire; Coherent, USA or Stradus 637-140) were used to emit circularly polarised laser radiation at 532 nm or 647 nm. Laser light was first reflected using a dichroic mirror (ZT405/488/532/640; Semrock, USA) before being directed to pass through an oil-immersion objective lens (CFI Apochromate TIRF Series 60× objective lens, numerical aperture = 1.49) to illuminate the sample. To ensure total internal reflection, the incident light was passed onto the sample at a critical angle of ∼ 67° for the glass/water interface present within the constructed flow cell. Fluorescent emission from AF555 and AF647 fluorophores was passed back through the objective lens and dichroic prior to being split by an additional T635lpxr-UF2 longpass dichroic mirror (Chroma, USA). Emission signal was then cleaned up further by passing through ET525/50 and ET690/50 m (Chroma, USA) filters before projection onto emCCD as two spatially split channels. The camera was operated at-70 °C for acquisition, with an electron multiplication gain of 900 and pixel distance of 160 nm. For experiments performed with the sample at 37 °C, temperature was maintained by an electronically heated flow-cell chamber attached to an objective heating jacket (Okolab, USA).

### Coverslip and flow cell assembly

To enable protein immobilisation, coverslips were first passivated as described previously^19^. Briefly, glass coverslips (24 × 24 mm) were cleaned by sonication in 100% ethanol and 5 M KOH twice (20 min each step) before aminosilanisation in 2% (v/v) 3-aminopropyl trimethoxysilane (Alfa Aesar, USA) for 15 min in acetone. Cleaned coverslips were then dried and incubated with 5 kDa NHS-ester methoxy-polyethylene glycol (mPEG) (150 mg/mL) and biotinylated-mPEG (bPEG; LaysanBio, USA) (6 mg/mL) prepared at a 25:1 ratio (mPEG: bPEG) as a ‘sandwich’ between two coverslips in a humidity chamber for a minimum of 4 h. PEGylated coverslips were then rinsed with milli-Q and subject to an additional PEGylation step as described above for an additional 20 h or overnight. PEGylated coverslips were then rinsed once more in milli-Q, dried with nitrogen gas and subsequently stored in a vacuum until use. Prior to use, PEGylated coverslips with incubated with neutravidin (0.2 mg/mL; BioLabs, USA) for 5 min. Coverslips were then rinsed, dried under nitrogen gas and fixed to a polydimethylsiloxane (PDMS) mould to construct the flow cell. Finally, to minimise non-specific binding of proteins to the coverslip surface, each microfluidic channel was incubated with 2% (v/v) Tween-20 for 30 min and then removed with imaging buffer (50 mM Tris [pH 7.5], 80 mM KCl, 10 mM MgCl_2_, 200 mM BME, 6 mM 6-hydroxy-2,5,7,8-tetramethylchroman-2-carboxylic acid [TROLOX]).

### Single-molecule fluorescence resonance energy transfer (smFRET) assay workflow

To monitor Fluc^IDS^ conformational changes as it is engaged by the Hsp70 machinery, fluorescently-labelled Fluc^IDS^ (50 pM final concentration) was introduced to the microfluidic channel and immobilised to neutravidin-functionalised coverslips. Excess, non-immobilised protein was then subsequently washed out of the flow cell with imaging buffer prior to imaging. To benchmark the folding sensor, native Fluc^IDS^ was imaged first in imaging buffer (Native) and following dilution out of 4 M GdHCl (Misfolded). Depending on the specific experiment, various combinations of HspA8, DnaJA2 and/or Hsp110 were introduced at the indicated concentrations for each experiment. ATP (5 mM) was present in all reactions unless stated otherwise.

### smFRET data analysis

Single-molecule trajectories were extracted from images and subsequently analysed using MASH-FRET software (version 1.2.2, accessible at https://rna-fretools.github.io/MASH-FRET)^20^. Briefly, the separated channels corresponding to donor and acceptor emission were aligned by a local weighted mean transformation generated by selecting colocalised spots in images of TetraSpack fluorescent beads. This transformation allowed for approximate FRET efficiency to be calculated through use of Equation 1 below:

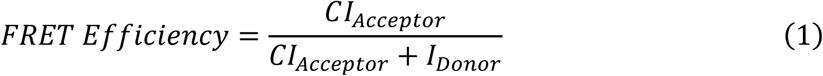

whereby CI_Acceptor_ represents the corrected acceptor fluorophore intensity given by *I_Acceptor_* – (g * *I_Donor_*) where *g* is the crosstalk correction factor defined as the ratio of fluorescence measured in both channels following excitation of a single-donor fluorophore. Minimal non-specific excitation of the acceptor fluorophore with the 532 nm donor laser was observed. FRET-competent molecules were selected by identifying traces with clear anti-correlation between donor and acceptor intensity with subsequent single-photobleaching events for both dyes. Traces were truncated at least 2 frames prior to photobleaching and passed through a denoising non-linear (NL) filter with the following parameter values selected: exponent factor for predictor weight – 5; running average window size – 1; factor for predicting average window sizes – 5. For downstream kinetic processing, the filtered traces were then fit with a Hidden Markov Model (HMM) using vbFRET (version vbFRET_nov12, accessible at https://sourceforge.net/projects/vbFRET/) to extract information about key state transitions and occupancy residence times.

### Kinetic analysis of smFRET data

HMM fits of FRET trajectories were further analysed to extract kinetic information about changes in client conformation during chaperone-mediated refolding. To distinguish FRET associated with the HspA8-induced conformational expansion, traces were designated as occupying either a *bound* (FRET efficiency < 0.3) or *unbound* (FRET efficiency > 0.3) state, with the time that a molecule resides in the bound state prior to entering the unbound state defined as *T_bound_* (and vice-versa for *T_unbound_*). As these residence times may be prematurely truncated due to dye photobleaching, the final occupied state in an individual trajectory was removed from the residence time dataset. Due to the smoothing effect applied during denoising, residence times shorter than the value described in equation 2 were also not considered, where NFA refers to the number of frames averaged during denoising.

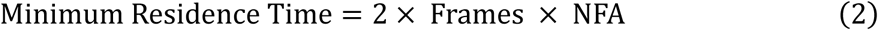

To determine kinetic rates underpinning transitions between *bound* and *unbound* states, the distribution of residence times were fit to either single-(equation 3) or double-exponentials (equation 4), where *T* refers to the distribution of residence times and Frac refers to the proportion of the function dictated by the faster of two kinetic rates, *k_1_*, such that both components sum to 1. Importantly, for double-exponential fitting, the kinetic rates were fixed such that the values of *k_1_* > *k_2_* for each fitting iteration in order to prevent swapping at initialisation.

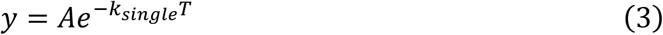

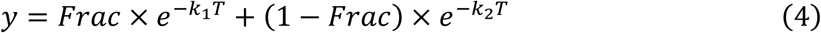

Fitting was performed by bootstrap (n = 500), where residence times were randomly sampled to equal the number of individual residence times within that treatment, fit to either a single-or double-exponential and the accuracy of the fit determined by plotting the residuals of both processes. The mean and standard error of *k_single_, k_1_, k_2_* and the value of Frac were extracted from the distribution of bootstrap results. Single-exponential fits were selected when higher order fits did not yield smaller residuals and were often characterised by a collapse of Frac in all iterations to exactly 0.5.

To characterise conformational changes during refolding, individual transitions could be visualised on transition density plots (TDPs), where the initial FRET state prior a transition was defined as ‘FRET before’, and the newly occupied state defined as ‘FRET after’. As transitions from a low FRET state (< 0.3) to a high FRET state (> 0.3) were identified to represent folding events from a HspA8-bound state, all such transitions were identified, aligned to the point of transition and the FRET efficiency 10 s pre-release and 50 s post-release were plotted as kernel density estimates (KDE) distributions.

### Unconstrained Gaussian fitting to identify native, misfolded and HspA8-bound states

To monitor the proportion of native, misfolded or HspA8-bound Fluc^IDS^ molecules during refolding, the FRET efficiency histograms were fit using an unconstrained mixed Gaussian model. Initial estimates of the peak mean were provided for each of the three states (∼ 0.1, 0.6 and 0.8 for HspA8-bound, native and compact misfolded, respectively) and the area under each curve were used to determine the contribution of each state to the total distribution.

### Statistical analysis and generation of figures

For comparisons of means between different treatments, data were analysed by either an unpaired t-test, or a one-way or two-way analysis of variance (ANOVA) followed by a Tukey’s multiple comparisons post-hoc test, whereby a value of *P* ≤ 0.05 was deemed to be statistically significant. All statistical analyses and figures presented were generated using GraphPad Prism 9 (GraphPad Software Inc; San Diego, USA) or custom-written Python scripts.

### Data, code, and materials availability

All code used for analysis and generation of figures found in the manuscript or the Supplementary Information can be accessed at the following Zenodo repository (https://doi.org/10.5281/zenodo.20389093; version 1.0.1). All ensemble and smFRET data can be accessed at the following Zenodo repository (https://doi.org/10.5281/zenodo.20473167; version 1.0.0). This study did not generate new materials. Further information and requests for any physical resources and reagents should be directed to the lead contact, H.E.

## Results

### The human Hsp70 chaperone system refolds chemically-denatured Fluc through active unfolding to conformationally-expanded states

To determine whether the most ubiquitous human Hsp70 chaperone system can refold clients via a similar mechanism to that described for the bacterial *E. coli* Hsp70 system^4–6^, we utilised a previously described firefly luciferase interdomain folding sensor (herein referred to as Fluc^IDS^) to observe changes in client conformation during chaperone-assisted refolding using smFRET^6^ (Figure 1A). First, we confirmed that the human Hsp70 system (consisting of HspA8 [Hsp70], DnaJA2 [Hsp40] and Hsp110 [NEF]) could refold chemically denatured Fluc^IDS^ to the native state using an enzymatic assay (Figure 1B). When Fluc^IDS^ was removed from denaturant in the absence of chaperones, negligible (< 5%) refolding to the native state was observed after 2 h (Figure 1B), consistent with previous work^6,7^. Likewise, refolding in the presence of both HspA8 and DnaJA2 did not result in a substantial return of enzymatic activity; however, addition of Hsp110 (which accelerates release of the client by HspA8) almost completely restored Fluc^IDS^ activity (∼ 90%) after 60 min of incubation (Figure 1B).

**Figure 1:**
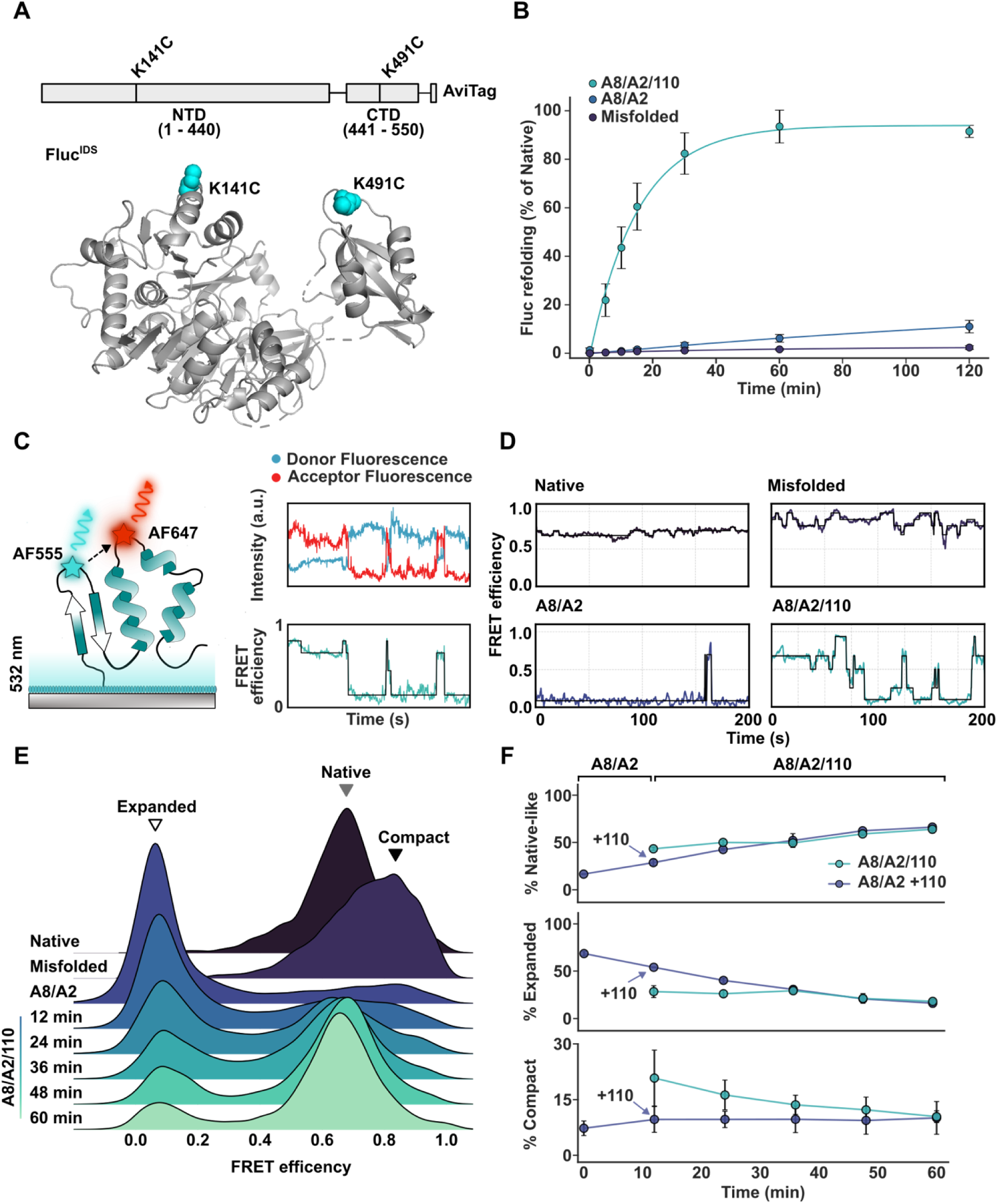
The complete human Hsp70 system resolves misfolded Fluc^IDS^ states through conformational expansion and promotes refolding events. **(A)** Schematic of Fluc crystal structure (PDB: 1LCI)^21^ with cysteine-substitutions for site-specific maleimide labelling indicated for Fluc^IDS^ (spheres). Fluc^IDS^ contains a C-terminal AviTag to enable immobilisation to a neutravidin-functionalised coverslip surface. **(B)** Fluc^IDS^ in 4 M GdHCl was diluted 100-fold to a final concentration of 10 nM into assay buffer (Fluc^IDS^ alone), in the presence of HspA8 (5 µM) and DnaJA2 (2 µM) alone (A8/A2), or supplemented with Hsp110 (0.5 µM; A8/A2/110). All conditions contained 5 mM ATP. Data shown depict mean ± SEM from 3 biological repeats. **(C)** Schematic depicting TIRF-based smFRET assays. AF555/AF647-labelled Fluc^IDS^ is immobilised to the neutravidin-functionalised coverslip surface and specifically excited at 532 nm. Fluorescence from both AF555 (donor) and AF647 (acceptor) fluorophores are monitored to calculate the FRET efficiency for individual molecules in real time. **(D)** Representative FRET efficiency traces for native Fluc^IDS^ (Native) or following dilution from 4 M GdHCl in the absence (Misfolded) or presence of HspA8 (3 µM) and DnaJA2 (2 µM) alone (A8/A2) or supplemented with Hsp110 (0.5 µM; + Hsp110). All conditions contained 5 mM ATP. Black line indicates the states determined by fitting raw FRET efficiency traces (coloured) with a Hidden Markov Model. Additional traces present in Figure S1B, Figure S3C. **(E)** Ridgeline plot depicting the FRET efficiency for Fluc^IDS^ molecules in their native state (Native), misfolded state (Misfolded) or in the presence of the indicated components of the Hsp70 system. Data collected over 60 min following initiation of refolding by addition of Hsp110. **(F)** Relative proportion of native-like, conformationally-expanded and compact misfolded molecules at different timepoints during refolding. Proportions were determined from the fit of a mixed Gaussian model with three states. Refolding was performed by addition of the complete Hsp70 machinery to misfolded Fluc^IDS1^ (A8/A2/110) or to HspA8-bound Fluc^IDS1^ (A8/A2 +110). The time at which Hsp110 was added is indicated (+ Hsp110). Refolding data are derived from 3 biological repeats and depicted as mean ± SEM. Data for all panels were derived and collated from a minimum of 315 molecules per condition.

To probe the mechanism by which Fluc refolding was achieved, we performed smFRET experiments with immobilised Fluc^IDS^ labelled with FRET-competent fluorophores (Figure 1A, C). This enabled the conformation of individual Fluc molecules to be monitored temporally during chaperone-mediated folding via changes in FRET efficiency (Figure 1C). Native Fluc^IDS^ exhibited stable FRET trajectories (Figure 1D; Figure S1) that were predominantly centred at 0.7 (Figure 1E). Chemical misfolding of the client was induced by addition of 4 M GdHCl to the flow cell, which resulted in Fluc^IDS^ occupying a low-FRET, expanded conformation (Figure S1A). Removal of denaturant with refolding buffer resulted in the collapse of Fluc^IDS^ to a broad, high-FRET distribution centred at ∼ 0.8-0.9, consistent with the formation of a compact chemically-misfolded state as observed previously^5,6^. Notably, the bulk luminescence assay confirms that this misfolded state represents an enzymatically inactive conformation of Fluc^IDS^ (Figure 1B).

Incubation of DnaJA2 and HspA8 with this misfolded state resulted in a shift to an ultra-low FRET state centred at ∼ 0.1 (Figure 1D-E), consistent with the client entering a conformationally-expanded HspA8-bound state comparable to that observed for the *E. coli* Hsp70, DnaK^6^. The persistence of the low-FRET HspA8-bound state explains the limited refolding observed in bulk refolding assays when incubated with DnaJA2 and HspA8 alone (Figure 1B), since stable client capture by HspA8 does not permit the client to sample native conformations. Crucially, incubation of native Fluc^IDS^ with DnaJA2 and HspA8 did not produce the same change in FRET efficiency (Figure S2A), indicating that these chaperones specifically recognise misfolded forms of Fluc.

Next, we supplemented the reaction with Hsp110, which resulted in Fluc^IDS^ returning to native-like FRET efficiencies over the course of 60 min (Figure 1E-F, *blue line*) consistent with the optimal refolding conditions observed in the enzyme activity assay when the complete Hsp70 system was present (Figure 1B). Similar results were observed when the complete Hsp70 system (*i.e*. DnaJA2, HspA8 and Hsp110) were added together to the misfolded Fluc (Figure 1F, *cyan line*), indicating that refolding kinetics are not substantially affected when refolding is initiated by the addition of Hsp110 to HspA8-bound Fluc^IDS1^; the only distinction is that the amount of misfolded Fluc^IDS^ remains lower.

The rate of transitions towards and away from the HspA8-bound low FRET state were accelerated in the presence of Hsp110 (Figure S2B), indicative of the NEF promoting faster HspA8 dissociation from Fluc. In instances where a molecule does not attain a stable native-like FRET efficiency, transitions back to HspA8-bound low FRET states can occur, which we define as a chaperone binding event. As the rate of these binding-and-release events is accelerated by Hsp110 (Figure 1D, Figure S2B), the client is provided additional opportunities for successful folding to occur. These data demonstrate that the human Hsp70 system can rapidly refold compact misfolded forms of Fluc^IDS^ to an enzymatically functional state via repeated cycles of HspA8-mediated client unfolding and release, comparable to that observed for the bacterial Hsp70 system^5,6^.

### Hsp110 promotes productive folding events from HspA8-bound states

Whilst Hsp70 acts to unfold misfolded client proteins through conformational expansion, one element of its role in client folding remains contentious – does Hsp70 play an active role in the *folding* of client proteins back to their native state or are clients left to independently refold upon release from Hsp70? To interrogate these two possibilities, we first investigated how Fluc^IDS^ spontaneously folds from a conformationally-expanded state (a context reminiscent of client release from the HspA8-bound state). First, immobilised Fluc^IDS^ was incubated with 4 M GdHCl to visualise transitions to an ultra-low FRET state (Figure 2A; Figure S1) comparable to the globally unfolded conformation Fluc^IDS^ adopts when bound by HspA8 (Figure 1E). Removal of denaturant resulted in clear transitions to the high-FRET, compact misfolded state (∼ 0.8 FRET efficiency; Figure 2A), with little to no intermediate states observed. Taking into account these key folding transitions - unfolding (*i.e.* transitions from FRET states greater than 0.5 to below 0.5) and folding (*i.e.* transitions from FRET states below 0.5 to above 0.5) (Figure 2A) - were time synchronised and plotted as 2D heatmaps (Figure 2B-D). These demonstrated that the refolding behaviour observed in individual traces are conserved at the ensemble level. By fitting the FRET efficiency distribution following attempted folding from denaturant (up to 10 s post-transition), the relative contributions from conformationally-expanded, native-like and compact-misfolded populations could be determined (Figure S3A). This demonstrated that ∼ 60% of molecules directly enter this compact misfolded state upon attempted folding from denaturant (Figure S3B). In contrast, folding events from the HspA8-bound state resulted in more native-like conformations, with FRET efficiencies centred at ∼ 0.7 in the presence of DnaJA2 and HspA8 (∼ 45% within 10 s of the folding event). Interestingly, when folding events were mediated by Hsp110, the proportion of native-like states were substantially higher (∼ 60%, Figure S3B). This effect was even more pronounced when the post-release period was extended to 50 s (Figure S4A-B), with ∼ 60% of molecules re-engaged by HspA8 and transition into the ultra-low, expanded conformation when incubated with HspA8 and DnaJA2 alone (Figure S4C), indicative of unsuccessful refolding. Occupancy in the ultra-low FRET state was substantially lower when Hsp110 was present (∼ 20%), indicating that refolding attempts are more successful. As such, Hsp110 plays a crucial role in the Hsp70-mediated refolding of Fluc^IDS^ by (i) promoting opportunities for folding by accelerating HspA8 dissociation, (ii) reducing collapse to compact misfolded states and (iii) coordinating folding events such that the folding trajectory of Fluc is biased towards the native state. Taken together, these results demonstrate that the human Hsp70 chaperone system guides Fluc^IDS^ down folding pathways that are distinct and more productive to attaining the native state than those that occur upon spontaneous collapse from a conformationally-expanded state.

**Figure 2:**
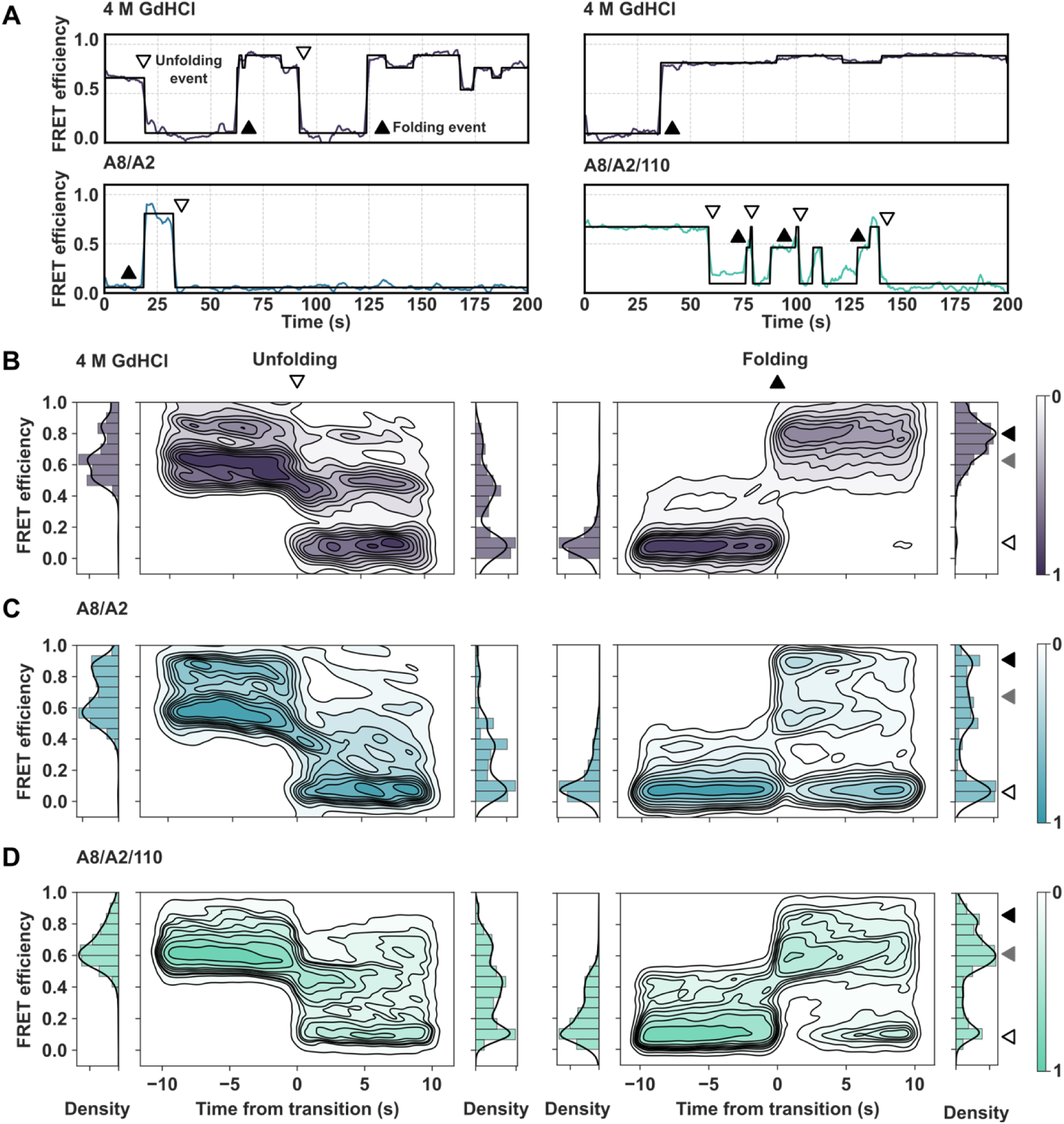
Folding events from HspA8-bound states biases Fluc^IDS^ towards the native state. **(A)** Example traces depicting Fluc^IDS^ intermittently supplemented with or diluted from 4 M GdHCl (top), or in the presence of A8/A2 (left) or A8/A2/110 (right). Transitions from above 0.5 to below 0.5 are defined as unfolding events and depicted by the white, downward triangle. Transitions from below 0.5 to above 0.5 are defined as folding events and denoted by a black, upward facing triangle. **(B)** Unfolding (left) and folding (right) events for Fluc^IDS^ supplemented with or following the removal of 4 M GdHCl corresponding to denaturant-induced unfolding and folding, respectively, in the presence of **(C)** A8/A2 alone or **(D)** with the complete Hsp70 system, A8/A2/110. White, grey and black filled triangles (right) indicate relative FRET efficiencies for the conformationally-expanded, native-like and compact misfolded states, respectively. FRET efficiency density 10 s prior and following a transition are depicted as histograms to the left and righthand side of the heatmap, respectively. Scale bar depicts relative density. Data for all panels derived from at least 174 molecules per condition.

We also investigated whether the conformational expansion of Fluc^IDS^ to low-FRET states by HspA8 and DnaJA2 alone (Figure 2C) differed compared to when Hsp110 was present (Figure 2D). For both conditions, transitions to the ultra-low FRET state (∼ 0.1) were most common. Interestingly, however, during conformational expansion by HspA8, Fluc^IDS^ adopted an intermediate FRET state centred at ∼ 0.4 before occupying the ultra-low FRET. Notably, this intermediate FRET state became more prominent when Hsp110 was present (Figure 2D). As this state does not appear in the absence of molecular chaperones (Figure 1E) but remains distinct to the ultra-low FRET conformation (∼ 0.1), this suggests the intermediate conformation may represent an alternatively bound or less saturated bound state of Fluc^IDS^. Additionally, a similar intermediate state at 0.4 FRET efficiency is also observed during the denaturant-induced unfolding of Fluc^IDS^ (Figure 2B), suggesting that this intermediate state may represent a structural local minimum for Fluc^IDS^ unfolding which is populated more frequently during HspA8-unfolding interactions. Regardless, these data demonstrate that HspA8 readily unfolds denatured Fluc^IDS^ to a predominantly ultra-low FRET state to allow refolding upon client release.

### The Hsp70 chaperone machinery protects against heat-induced inactivation of Fluc^IDS^ via a partial unfolding mechanism

Having visualised the mechanism by which HspA8 actively refolds a globally denatured protein, we next sought to investigate whether this mechanism was also employed for maintenance of native states during heat stress. First, we performed ensemble-based luminescence assays to confirm that Fluc^IDS^ misfolds and becomes non-functional when exposed to mild heat stress (Figure 3A). When native Fluc^IDS^ was incubated at 37 °C, the measured enzyme activity decreased over time such that it exhibited only ∼ 25% of the native activity after 2 h. In contrast, incubation with the complete Hsp70 system during mild heat stress significantly protected Fluc^IDS^ against misfolding, with ∼ 70% of the native signal remaining intact. Interestingly, removal of the heat stress after 2 h of incubation did not result in a meaningful increase in Fluc^IDS^ activity in either the absence or presence of the Hsp70 chaperones. These data suggest that the Hsp70 system recognises heat denatured Fluc^IDS^ and acts to maintain a predominately folded and functional population during heat stress. The inability of the Hsp70 system to maintain a completely folded client protein population during heat stress has been previously reported^17,21^: this non-native fraction is thought to likely represent the formation of small aggregates that cannot be recovered by Hsp70. To remove aggregation as a possibility, we performed comparable experiments at 37 °C using our smFRET assay in which Fluc^IDS^ is spatially resolved by immobilisation to the coverslip.

**Figure 3:**
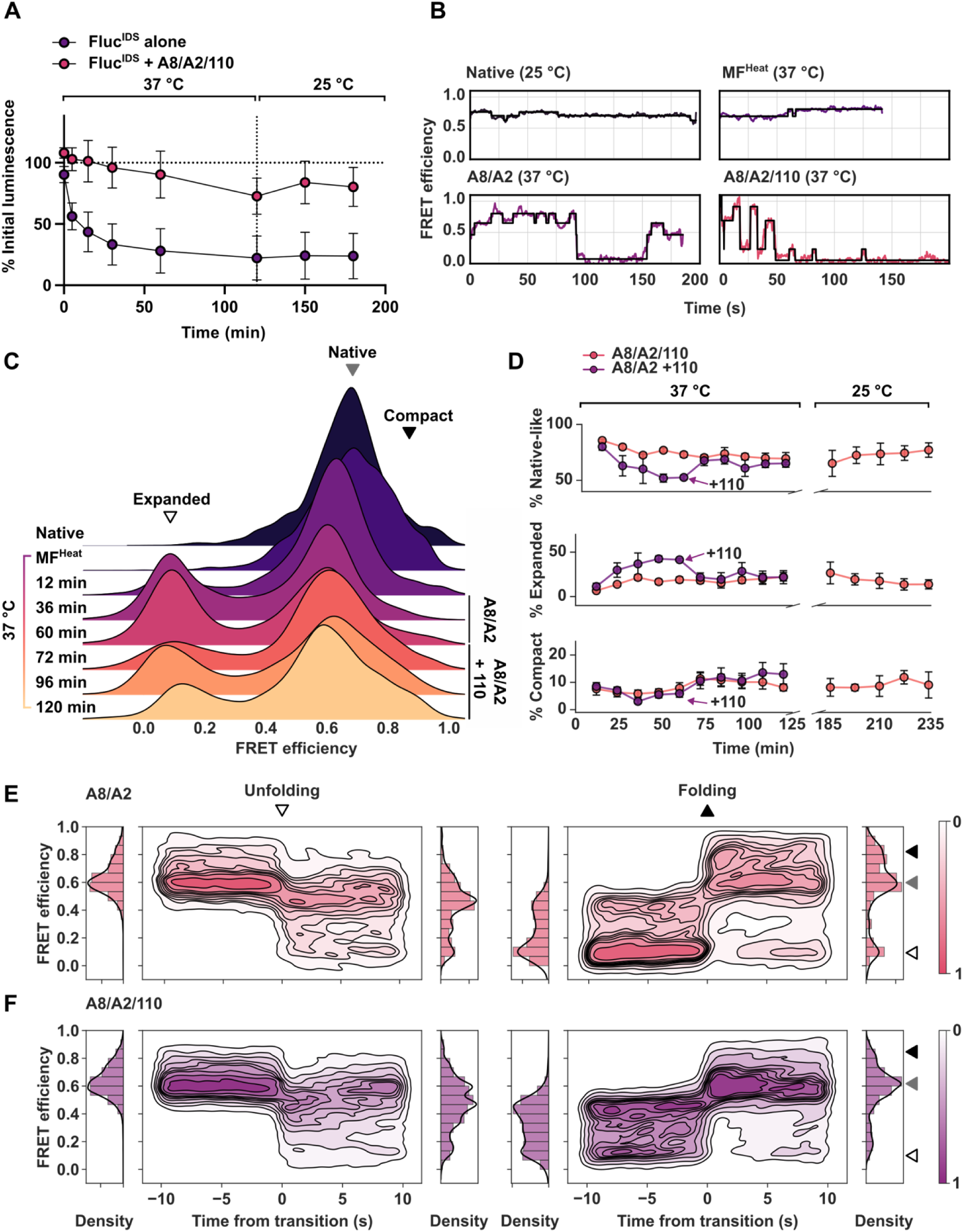
HspA8 maintains the native state of Fluc^IDS^ under conditions of heat stress by partially unfolding non-native regions. **(A)** Bulk enzymatic assay whereby Fluc^IDS^ was incubated at 37 °C alone or in the presence of the complete Hsp70 system (3 µM HspA8, 2 µM DnaJA2, 0.5 µM Hsp110) over 2 h, with luminescence normalised to the initial native signal. After 2 h, both treatments were returned to room temp (25 °C) and monitored for a further 60 min. Data are representative of 3 biological repeats and depicted as mean ± SEM. **(B)** Representative smFRET traces of Fluc^IDS^ alone incubated at room temperature (Native (25 °C)) and during heat stress (MF^Heat^ (37 °C)). Traces are also provided depicting Fluc^IDS^ at 37 °C in the presence of 3 µM HspA8 and 2 µM DnaJA2 (A8/A2 (37 °C), or in the presence of the complete Hsp70 system (A8/A2/110 (37 °C)). **(C)** Ridgeline plot depicting the FRET efficiency of Fluc^IDS^ when incubated under native conditions at 25 °C (Native), following exposure to heat stress at 37 °C for 1 h in the absence (MF^Heat^ (60 min)) and presence of A8/A2 alone (A8/A2), or supplemented with Hsp110 (A8/A2 + 110). Data are presented as mean ± SEM from 3 biological repeats. **(D)** Relative contributions of native-like, conformationally-expanded and compact misfolded states of Fluc^IDS^ at different timepoints during heat stress (37 °C) and upon return to room temperature (25 °C) as determined by fitting the FRET efficiency distribution with the sum of 3 Gaussians. **(E)** 2D heatmaps depicting unfolding (left) or folding (right) events for Fluc^IDS^ incubated at 37 °C in the presence of A8/A2 alone or supplemented with the complete chaperone system (A8/A2/110). **(F)** Relative density is depicted by the scalebar. The FRET efficiencies representing the expanded, native-like and compact states are indicated by the white, grey and black triangles, respectively. Data for all panels were derived from at least 173 molecules per condition.

Interestingly, prolonged incubation of Fluc^IDS^ at temperatures whereby misfolding and loss of activity occurs in the bulk enzyme assay (Figure 3A) did not substantially alter the FRET efficiency distribution compared to the native state measured without heat stress (*i.e*., at 25 °C; ∼ 0.7) (Figure 3B-C). Indeed, individual FRET trajectories for Fluc^IDS^ heated at 37 °C did not demonstrate significant changes in FRET efficiency (*i.e*. no clear transitions to high-or low-FRET states indicative of heat-induced global misfolding or unfolding, respectively) (Figure 3B). However, incubation of heat stressed Fluc^IDS^ with HspA8 and DnaJA2 resulted in transitions to low-FRET states (Figure 3B) that accumulated in abundance over time (Figure 3C) such that, after 1 h of incubation, ∼ 40% of the FRET signal was from molecules in a low FRET state. Since HspA8 does not bind to native Fluc^IDS^ (Figure S2A), these data indicate that Fluc^IDS^ does indeed misfold under the heat stress conditions used in these experiments; however, this misfolding occurs in such a way that it does not result in large conformational changes such as that which occurs during the formation of a compact misfolded state with high-FRET. Thus, the heat-denatured misfolded state of Fluc (at native-like FRET efficiencies) appears to be distinct to the high-FRET compact misfolded state generated using chemical denaturants (> 0.8, Figure 1E), possibly since heat stress results in (i) minor structural rearrangements not detectable by FRET or (ii) misfolding occurring in other regions of the protein not probed by the Fluc^IDS^ FRET sensor. Interestingly, transitions from a low-FRET (∼ 0.1) to high-FRET state (> 0.8) were observed, albeit rarely, when heat stressed Fluc was incubated with HspA8 and DnaJA2 (Figure 3B, F; Figure S5). Since these high-FRET states do not frequently appear in the absence of these chaperones, this suggests that HspA8-induced conformational expansion of Fluc increases the accessible folding landscape and enables access to alternative folding and misfolding states that are not likely to be sampled under heat stress alone.

To quantify the relative proportions of molecules occupying conformationally-expanded low-FRET, native-like or compact-misfolded high-FRET conformations, the FRET efficiency distributions at different timepoints were determined (Figure S2; Figure 3D). When Fluc^IDS^ was incubated in the presence of HspA8 and DnaJA2 alone, the abundance of the low-FRET state increased over time (from ∼ 5% to ∼ 40% after 1 hour) and was accompanied by a corresponding decrease in the proportion of molecules exhibiting a native-like FRET efficiency of ∼ 0.7 (from 80% to 50% after 1 hour) (Figure 3C-D). Addition of Hsp110 at this point resulted in a return to native-like FRET distributions (∼ 70% at 25 min after addition of Hsp110; Figure 3C-D) and, crucially, the proportion of Fluc^IDS^ in a low-FRET state decreased to ∼ 10%, indicating that HspA8 did not re-engage with these molecules and that productive folding to the native state had occurred (Figure 3D). Interestingly, addition of Hsp110 also increased the formation of high-FRET (> 0.8 FRET) species reminiscent of the chemically-induced compact misfolded state (Figure 1E), which increased in abundance to ∼ 18% before plateauing (Figure 3D). Delayed addition of the NEF to heat-stressed Fluc^IDS^ that had been incubated for 60 min with HspA8/DnaJA2 resulted in the same occupancy of all three Fluc^IDS^ states (*i.e.* conformationally-expanded, native-like and compact misfolded) as those observed when the complete Hsp70 system was present at initiation of heat-stress. Thus, in the absence of a NEF, HspA8 acts primarily in a *holdase* capacity and does not alter the proportion of molecules present in a folding-competent state. The complete Hsp70 chaperone system maintains Fluc^IDS^ in an equilibrium of states such that ∼ 70% of the molecules remain in a native-like state (Figure 3D); however, this equilibrium is maintained by active unfolding and refolding of these molecules. When this system was returned to 25 °C, the proportion of molecules in the native state increased to ∼ 80% such that it was comparable to the population present at the start of the incubation (Figure 3D). These results are therefore consistent with the negligeable increase in enzyme activity observed in the bulk luminescence assay when the system was returned to 25 °C (Figure 3A). As aggregation of Fluc cannot occur under the smFRET experimental conditions used in this work (since individual molecules are spatially separated through immobilisation to the coverslip surface), this non-native fraction likely represents molecules that have yet to be refolded and continue to undergo conformational remodelling by HspA8 (Figure 3B).

We next investigated whether the mechanism by which Hsp70 remodels heat stressed Fluc^IDS^ differs from that observed for chemically misfolded Fluc^IDS^. To do this, we first identified HspA8 binding events, defined as FRET transitions to a conformationally-expanded state (FRET efficiency < 0.5) from a more compact state (> 0.5, Figure 3E-F). Interestingly, the majority of HspA8 binding events resulted in only partial Fluc^IDS^ unfolding in both the absence (Figure 3E) and presence of Hsp110 (Figure 3F), as demonstrated by high FRET efficiency density at ∼ 0.4 compared to the typical HspA8-bound, ultra-low FRET state (∼ 0.1). Notably, this intermediate conformation is rarely populated in the FRET distributions (Figure 3C), indicating that it is a short-lived transition state that is masked at steady state by the more stable, fully expanded conformation. As transitions to these intermediate states were frequently observed, it appears that only partial expansion of Fluc^IDS^ is required to resolve heat-induced misfolded states to promote folding. This differs substantially to the way in which Hsp70 acts to resolve the compact, denaturant-induced misfolded Fluc^IDS^ state (Figure 1E), whereby HspA8 binding rarely results in the intermediate FRET state (∼ 0.4) (Figure 2D-E) or bypasses it entirely en route to the ultra-low FRET expanded state (∼ 0.1).

Interestingly, many folding events were initiated from the partially expanded intermediate, particularly in the presence of Hsp110 (Figure 3E-F). This directly contrasts results from when chemically-misfolded Fluc^IDS^ was refolded at room temperature, where folding events originated primarily from the completely expanded state (Figure 2C-D). Given that Hsp110 promotes dissociation of HspA8 from the client, the increased occupancy of this intermediate state provides further evidence that it may represent an alternately bound HspA8-Fluc^IDS^ complex generated via decreased HspA8 binding. Folding events initiated from the partially expanded state (Figure S6A, C) also resulted in substantially more native-like conformations than those from the fully expanded, ultra-low FRET state (Figure S5B, D-E); notably, this observation occurred irrespective of the presence of Hsp110. Thus, these findings suggest that partial unfolding of Fluc^IDS^ by HspA8 is sufficient to facilitate its refolding to the native state during heat stress. In fact, complete expansion of Fluc^IDS^ by HspA8 under heat stress conditions may permit alternate misfolded states to accumulate following non-productive folding. Collectively, these data demonstrate that HspA8 preserves a native Fluc^IDS^ population under heat stress by actively remodelling the client conformation through an unfolding mechanism analogous to that used during refolding of chemically-denatured clients. However, the degree of expansion required to facilitate folding to the native state is less extensive than that necessary to refold a globally misfolded client.

### The capacity of Hsp70 to unfold Fluc at elevated temperature is insufficient for resolving denaturant-induced misfolding

As HspA8 exhibited distinct unfolding behaviour toward Fluc^IDS^ at room temperature (25 °C) and 37 °C, we sought to determine whether these differences were observed due to a change in temperature alone, the nature of the misfolded client, or a combination of both. To do this, we assessed the ability of the human Hsp70 system to refold two distinct Fluc^IDS^ misfolded states at 37 °C; those generated by chemical denaturation in 4 M GdHCl (chemically misfolded, herein referred to as MF^Chem^) or by incubation at 37 °C for 1 h (heat misfolded, herein referred to as MF^Heat^). Interestingly, the complete Hsp70 system could only restore MF^Chem^ activity to ∼ 15% of the native luminescence (Figure S7A), which is markedly lower than the near complete reactivation observed at room temperature (∼ 85%, see Figure 1B). Notably, the same combination of chaperones was able to mostly protect a native Fluc^IDS^ population from misfolding at identical temperatures (Figure 3C-D). To interrogate why refolding of MF^Chem^ was impaired at elevated temperatures, we exploited our smFRET approach.

Dilution out of denaturant at 37 °C caused Fluc^IDS^ to adopt the characteristic high-FRET misfolded conformation (MFChem; Figure S7B) that was also observed at room temperature. However, addition of HspA8 and DnaJA2 to this conformation resulted in only a minor shift to the ultra-low FRET state (∼ 0.1) (Figure 4A; Figure S7B). Inspection of the unfolding trajectories of individual molecules revealed that this ultra-low FRET state was only transiently occupied before molecules returned either to the partially unfolded intermediate (FRET ∼ 0.4) or collapsed back into a compact misfolded state (FRET ∼ 0.8) (Figure S7C-D). Indeed, time spent in unfolded conformations (*i.e.* FRET < 0.5) before transition to more compact states was significantly shorter for MF^Chem^ incubated in the presence of A8/A2 at 37 °C (*T_bound_* ≈ 20 s) than at room temperature (*T_bound_* ≈ 50 s) (Figure 4B). Additionally, the average FRET state occupied immediately prior to a folding event (*i.e.* transition from < 0.5 FRET to > 0.5 FRET) was significantly lower for MF^Chem^ incubated with A8/A2 at 25 °C (FRET efficiency ∼ 0.15) compared to the release point at 37 °C (∼ 0.23; Figure 4C). Therefore, not only was the complex formed between MF^Chem^ and HspA8 less stable as indicated by shorter *T_bound_* times, but the degree of unfolding imparted on the client by HspA8 decreased under heated conditions. Addition of Hsp110 to this partially expanded client-HspA8 complex failed to promote complete recovery of native-like conformations (Figure S7B), consistent with the minimal return in enzymatic activity observed in the bulk assay (Figure S7A). These data demonstrate that elevated temperature alone negatively impacts the capacity of HspA8 to fully resolve MF^Chem^, suggesting altered binding and release kinetics that ultimately disfavour complete unfolding and therefore refolding of certain clients at higher temperatures.

**Figure 4:**
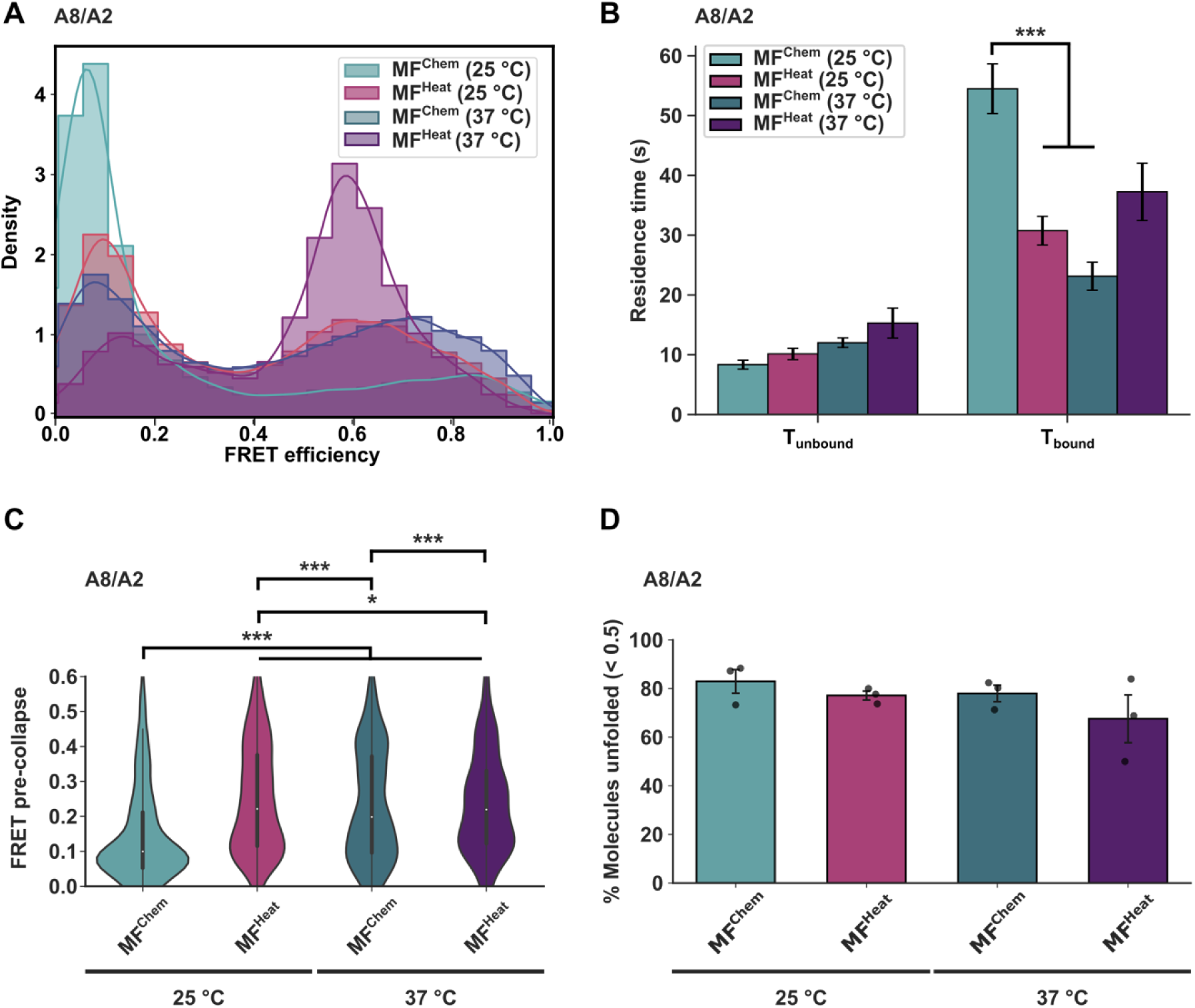
Partial expansion of Fluc^IDS^ by HspA8 at elevated temperatures is insufficient for refolding of the chemically-misfolded protein to the native state. **(A)** Ridgeline plot depicting smFRET efficiency of misfolded Fluc^IDS^ via incubation with 4 M GdHCl (MF^Chem^) or heat for 1 h at 37 °C (MF^Heat^) incubated with 3 µM HspA8 and 2 µM DnaJA2 at either 25 °C or 37 °C. **(B)** Mean residence time above (*T_unbound_*) or below (*T_bound_*) a threshold of FRET 0.5 for MF^Chem^ and MF^Heat^ incubated with A8/A2 for 30 min at either 25 °C or 37 °C. Data are presented as mean ± SEM from 3 biological repeats, with a minimum of 73 events per treatment. A one-way ANOVA followed by Tukey’s HSD post-hoc test was performed, where *** indicates significance (*P* < 0.001). Comparisons not indicated were not significant (*P*> 0.05). **(C)** FRET efficiency occupied immediately prior to transition to a collapsed state (FRET > 0.5) for MF^Chem^ and MF^Heat^ at 25 °C or 37 °C. Data are representative of a minimum of 37 dwell times per treatment. Means were assessed by two-way ANOVA, followed by Tukey’s HSD post-hoc test, where *** and * indicate significance with *P* values of less than 0.001 and 0.05, respectively. **(D)** Proportion of molecules that visit an A8-mediated unfolded state (FRET < 0.5) at any point during imaging. Data are presented as mean ± SEM from 3 biological repeats. A two-way ANOVA was performed with no statistically significant differences between groups determined. Data for all panels were derived from a minimum of 260 molecules per treatment.

It remains unclear whether intrinsic differences in the conformation of the misfolded client itself also influence the ability of HspA8 to bind to and unfold the client. To investigate this, we incubated either MF^Chem^ or MF^Heat^ at room temperature (25 °C) with HspA8 and DnaJA2 and observed the ability of HspA8 to generate the stable conformationally-expanded state (Figure 4A) and assessed the kinetics associated with entering this state (Figure 4D). We first identified molecules entering partially unfolded conformations (*i.e.* visiting FRET efficiencies < 0.5) as these states are only occupied when HspA8 interacts with misfolded Fluc^IDS^ (Figure S2A). The time taken for binding and subsequent unfolding to a partially unfolded state for both misfolded clients were similar (*T_unbound_* ≈ 10 s), suggesting both conformers were recognised and bound by HspA8 (Figure 4B); however, the HspA8-mediated unfolded state was significantly longer and more stable for MF^Chem^ (*T_bound_* ≈ 50 s) than MF^Heat^ (≈ 30 s) (Figure 4B). Strikingly, this relationship was inverted at elevated temperatures, with MF^Heat^ bound for slightly longer (*T_bound_* ≈ 35 s) compared to MF^Chem^; (*T_bound_* ≈ 20 s) (Figure 4B), although this did not reach statistical significance. Additionally, refolding events were initiated from significantly lower FRET efficiencies for MF^Chem^ (∼ 0.15) compared to MF^Heat^ (∼ 0.25) at room temperature (Figure 4C), consistent with the decreased expansion of the client observed during refolding at 37 °C (Figure 3E-F). Indeed, whilst 80% of molecules for both conformers were found to visit the partially unfolded state when in the presence of A8/A2 (*i.e.* experienced transitions to FRET values < 0.5) regardless of temperature (Figure 4D), MF^Heat^ was found to have a significantly lower proportion enter the ultra-low FRET state (*i.e.* transitions to FRET values < 0.3) at 37 °C (∼ 20%; Figure S8A) in comparison to MF^Chem^ (∼ 60%). This result also coincides with MF^Heat^ spending significantly more time in FRET states > 0.3 (*T_unbound_* ≈ 18 s) than MF^Chem^ at 37 °C (*T_unbound_* ≈ 10 s; Figure S8B). Collectively, these data describe a system in which the distinct misfolded conformations of the same client protein possess intrinsic structural differences that influence how they are remodelled by HspA8.

### The stability of the HspA8-Fluc^IDS^ complex governs the switch in chaperone mechanism from holding to folding

Productive folding of a client by the Hsp70 machinery reflects a balance between the need for efficient resolution of misfolded structures through active unfolding and NEF-mediated client release that permits the formation of native contacts required for function. Indeed, under conditions of excessive Hsp70 binding, clients become trapped in an expanded conformation and folding is inhibited^22^. To better understand how HspA8 maintains this equilibrium, we quantified the kinetics of folding and unfolding of MF^Chem^ at room temperature. As expected, the time required for HspA8 to bind and unfold Fluc^IDS^ was independent of Hsp110 (*T_unbound_* ≈ 10 s) (Figure 5A). In contrast, the lifetime of the low FRET HspA8-bound state was quite stable in the absence of NEF (*T_bound_* ≈ 50 s) and substantially shorter when Hsp110 was present (≈ 20 s) (Figure 5A). Thus, Hsp110 accelerates escape from the HspA8-bound conformation and increases the frequency with which MF^Chem^ can attempt refolding to the native state (Figure S2B). Interestingly, the distribution of dwell times for *T_bound_* were poorly described by a single-order process when MF^Chem^ was incubated with HspA8 and DnaJA2 alone (Figure S9A), suggesting that additional kinetic features regulate the longevity of the unfolded state. This was not the case for *T_unbound_*, which was clearly defined by a single kinetic rate (Figure S10). Indeed, a bi-exponential model best fit the *T_bound_* data, with the kinetics of release from the HspA8-bound state dictated by a fast (herein referred to as *k_fast_*) and a slow (referred to as *k_slow_*) rate (Figure 5B,G; Figure S9B). In the presence of HspA8 and DnaJA2 alone, approximately 40% of the HspA8-bound residence times were governed by *k_fast_* (0.1 s^-1^; Figure 5B-C), which was ∼ 10-fold faster than *k_slow_* (0.01 s^-1^; Figure 5B). Importantly, *k_slow_* was not observed when Hsp110 was present (Figure S9A,C), suggesting that NEF-mediated dissociation of HspA8 from Fluc^IDS^ prevents formation of a stable HspA8-MF^Chem^ complex. In contrast, *T_unbound_* was well described by a single-order unfolding rate (*k_unfold_* ≈ 0.15 s^-1^) that was independent of Hsp110 (Figure 5B).

**Figure 5:**
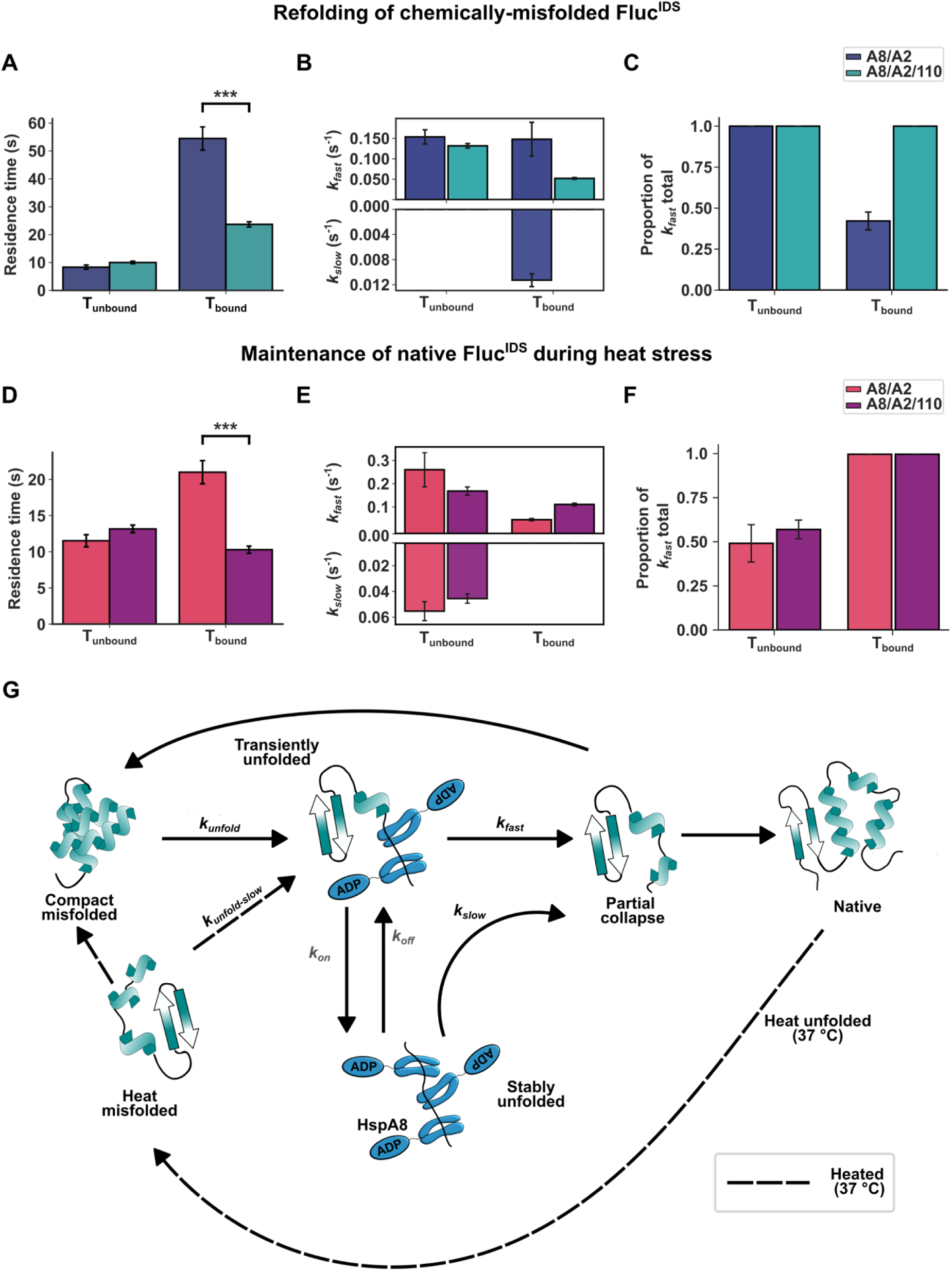
Hsp110 destabilises HspA8-Fluc complexes to promote opportunities for folding. **(A)** Residence time above (*T_unbound_*) or below (*T_bound_*) a threshold of 0.5 for MF^Chem^ in the absence (A8/A2) or presence of Hsp110 (A8/A2/110). Data are representative of a minimum of 246 residence times depicted as mean ± SEM from 3 biological repeats. **(B)** Kinetic rate constants for *k_fast_* (top) and *k_slow_* (bottom) underpinning mean dwell times for bound and unbound states. Data are representative of mean ± SEM of 500 bootstrap fits. Treatments with a single bar are indicative of kinetic processes with only one kinetic rate. **(C)** Proportion of kinetic fit dictated by the fast rate, *k_fast_*. **(D)** Residence times for MF^Heat^ as described in (A). Data are representative of a minimum of 133 residence times depicted as mean ± SEM pooled from 3 biological repeats. **(E)** Kinetic rate constants for MF^Heat^ as described in (B). **(F)** Proportion of kinetic fit dictated by *k_fast_*. **(G)** Schematic illustrating the proposed pathways underlying the chaperone action of HspA8. Compact misfolded clients are bound and unfolded by HspA8 with a single rate, *k_unfold_*. Two bound states are then possible – a shorter-lived HspA8-client complex with occupancy described by *k_fast_* and a more stable unfolded state represented by *k_slow_*. The longer-lived HspA8-bound state is characterised by association (k_on_) and dissociation (*k_off_*) of additional HspA8 subunits. Dissociation of HspA8 below a minimum stoichiometry threshold results in collapse to a partially folded intermediate, which may then fold into the native state or be re-engaged with HspA8. Natively folded clients may partially misfolded (e.g. due to heat stress) and are subsequently recognised and partially unfolded by HspA8 with a distinct rate, *k_unfold,slow_*. In some cases, the binding of additional HspA8 molecules leads to conformational expansion of the client. Release from this HspA8-bound conformationally-expanded state may result in misfolding to a compact misfolded state. A one-way ANOVA followed by Tukey’s HSD post-hoc test was performed for data presented in (A) and (D), where *** indicates significance (*P* < 0.001). Comparisons not indicated were not significant (*P*> 0.05).

As dissociation of a single Hsp70 monomer from a client follows single-order kinetics^23,24^, these findings suggest that the stability of the HspA8-bound state is additionally regulated by non-kinetic and/or physical features of the system. There are ∼ 10 predicted Hsp70 binding sites present on Fluc^25^; as such, it is likely that the low-FRET population represents an ensemble of HspA8-Fluc complexes with varying stoichiometries and binding topologies, defined by multiple on/off rates. Indeed, proposed mechanisms of Hsp70 chaperone function, such as entropic pulling^8^ and volume exclusion^4^, rely on the coordinated action of multiple Hsp70 molecules to drive unfolding of the client. Thus, the two kinetic rates governing occupancy of the HspA8-bound state of Fluc^IDS^ (*i.e*., *k_fast_* and *k_slow_*) may reflect differences in the binding saturation of HspA8 on a single Fluc^IDS^ molecule. Short-lived unfolded states, dictated by *k_fast_*, likely reflect the dissociation kinetics of a single HspA8 from a HspA8-bound state in which a single dissociation event is sufficient to allow folding (Figure 5G). In contrast, the slower rate (*k_slow_*) may reflect HspA8-bound states above this threshold in which many more successive HspA8 dissociation events are required to permit Fluc compaction/folding. Critically, both rates are described by similar FRET efficiency distributions as FRET is unable to report on further conformational expansion of the client and additional HspA8 binding does not appreciably further decrease FRET. When the balance between association and dissociation of HspA8 is tipped towards association (as occurs in the absence of Hsp110), the unfolded conformation is additionally stabilised (as observed in Figure 5A-B, A8/A2). Shifting this equilibrium to favour dissociation by addition of Hsp110 results in elimination of this more stable kinetic feature represented by *k_slow_* (Figure 5A-B, A8/A2/110). Together, these results demonstrate that HspA8 binds chemically-misfolded Fluc^IDS^ and maintains it in a conformationally-expanded ensemble with distinct stabilities. By increasing HspA8 dissociation, Hsp110 promotes more frequent opportunities for productive folding and prevents client entrapment in a long-lived, non-folding-productive, HspA8-bound state.

### Maintenance of the native state of heat stressed Fluc^IDS^ by Hsp70 is characterised by short-lived client-chaperone interactions

Given that HspA8 maintained heat stressed Fluc^IDS^ in a native-like conformation via a mechanism distinct from that used to refold a globally denatured state, we investigated whether the underlying kinetics were similarly distinct. Similar to the kinetics for MF^Chem^, the time to unfold MF^Heat^ was also independent of Hsp110, with a value of *T_unbound_* ≈ 20 s for both treatments (Fig 5D). Interestingly however, dwell time distributions for *T_unbound_* under heat stress were best described by a bi-exponential model, both with and without Hsp110 (Figure 5E-F; rationale described in Fig S9-S10). This added kinetic complexity likely reflects fundamental differences between the refolding mechanism of HspA8 for chemically-misfolded Fluc^IDS^ and protection against heat-induced denaturation. In chemically denatured samples, the entire population begins in a misfolded state, enabling immediate HspA8 engagement and unfolding which is governed by a single rate constant (*k_unfold_).* In contrast, under heat stress conditions, the client molecule population is initially, predominantly in the native state, and HspA8 can only bind client molecules as they spontaneously misfold. Thus, the slower unfolding rate (*k_unfold,slow_*) detected at 37 °C may represent heterogeneity in HspA8 binding events to exposed hydrophobic regions during early misfolding before large-scale conformational expansion can occur (Figure 5G). Additionally, these early HspA8 binding events may also be particularly sensitive to Hsp110-mediated dissociation. This sensitivity may arise from limited HspA8 binding site availability in early misfolding intermediates, such that NEF-driven dissociation occurs before sufficient HspA8 accumulation can promote unfolding. As expected, Hsp110 substantially decreases the stability of the HspA8-Fluc complex, with *T_bound_* observed to be significantly reduced (from ∼ 20 s to 10 s) upon addition of the NEF (Figure 5D). These relatively brief excursions to the HspA8-bound state both with and without Hsp110, coincides with the absence of *k_slow_* for the *T_bound_* state of M^Heat^, in contrast to MF^Chem^ (Figure 5E; rationale described in Fig S9-S10). Collectively, these results indicate that HspA8 maintains the native Fluc^IDS^ state during heat stress through transient and minor conformational expansion events to facilitate rapid refolding (Figure 5G). In this context, HspA8 functions not as a holding chaperone but as a dynamic surveillance factor, engaging locally misfolded clients briefly to prevent irreversible misfolding while preserving access to the native state.

## Discussion

Hsp70 is frequently described as the ‘central hub’ of the chaperone network, reflecting its broad role in maintaining proteostasis through protein quality control and protection against cellular stress. Central to this role is its ability to both remodel misfolded proteins into functional states and preserve native proteins under destabilising conditions. Despite extensive study, the mechanistic basis by which the constitutively expressed human Hsp70, HspA8, performs these functions has remained incompletely defined. Here, we directly visualised ATP-dependent refolding and native-state maintenance of a single client protein (Fluc) by the human Hsp70 machinery and reveal how modulation of client-chaperone complex stability governs the switch between *holdase* and *foldase* function.

We show that HspA8 binding induces rapid interdomain expansion to resolve misfolded Fluc states and maintains the client in an ensemble of unfolded conformations similar to that observed for the bacterial Hsp70 homolog, DnaK^4–7,21,26^; this underscores the evolutionary conservation of Hsp70-mediated client unfolding as a fundamental mechanism to refold proteins to their native state. We further demonstrate that Hsp110 acts as a molecular switch that promotes HspA8 dissociation from the client to enable productive refolding. In this way, Hsp110 converts HspA8 from a *holdase* chaperone that stabilises unfolded clients into a *foldase* that facilitates refolding to the native state.

There is growing evidence that Hsp70 does not only resolve misfolded clients but can actively direct folding trajectories toward native states^5,21,27–29^. Consistent with this view, we find that refolding of Fluc from conformationally-expanded states is mechanistically distinct when originating from HspA8-bound states compared to release initiated from chemical denaturant, suggesting that the presence of HspA8 plays a critical role in directing proteins down productive folding pathways. Strikingly, productive folding events are enhanced by Hsp110 (Figure S4C-D). This may be due to Hsp110 coordinating the dissociation of HspA8 subunits to favour correct folding or by increasing the cycling of binding-and-release events such that misfolding proteins are ‘captured’ prior to collapsing completely to the compact misfolded state. Moreover, as Fluc contains ∼10 predicted Hsp70 binding sites^25,30,31^, asynchronous release of individual HspA8 molecules likely permits localised refolding of the released region while adjacent domains remain shielded by bound chaperones^27,28^. Such a mechanism would suppress non-native long-range interactions between distant regions of the polypeptide, effectively smoothing the folding energy landscape, accelerating folding kinetics and increasing the overall probability of successful refolding.

Although it is well established that Hsp70 can maintain a native protein population under conditions of heat stress^5,15,21,32,33^, the extent to which this protection reflects explicit *holdase* or *foldase* chaperone function has remained unclear. Previous studies suggest that DnaK can bind and stabilise native conformations of certain clients^15,34^, whereas others report active unfolding of heat-misfolded Fluc^5^. Our data reveal that at elevated temperature (37 °C), HspA8 and DnaJA2 rapidly detect and unfold heat stressed Fluc^IDS^ via a mechanism similar to that used to refold the protein from the compact state formed following chemical denaturation. However, in the presence of Hsp110, these unfolding events are much shorter-lived and often result in only partial expansion of the client. Because Fluc^IDS^ remains predominantly functional in the presence of the complete Hsp70 chaperone system at elevated temperatures, transient unfolding to partially expanded states appears sufficient to preserve the native state, where global unfolding of the protein is not required.

Folding from more extensively expanded states at 37 °C increases the formation of compact high-FRET states reminiscent of the misfolded ensemble generated by chemical-denaturation (*i.e.* MF^Chem^). These compact misfolded states do not always appear during heat-stress and interestingly, require more extensive HspA8 engagement to refold (Figure 4A-B). This suggests that unsuccessful chaperone engagement may produce alternatively misfolded structures that may not otherwise be present, which imposes a greater energetic and chaperone burden to resolve these states. Moreover, we demonstrate that HspA8 binds to chemically (MF^Chem^) and thermally (MF^Heat^) misfolded conformers of Fluc with distinct client-chaperone kinetics and refolding requirements. This distinction likely reflects fundamental differences between the two misfolded conformers: MF^Chem^ arises from near-complete loss of structure, whereas heat stress induces more localised misfolding in MF^Heat^. Consistent with this, recent molecular dynamic simulations indicate that Fluc heated to 42 °C undergoes limited, region-specific misfolding while maintaining much of its native state structure^35^. The modest degree of unfolding by HspA8 required to maintain Fluc^IDS^ in a native state aligns with these findings. Such differences in how Hsp70 interfaces with distinct misfolded conformations of the same client protein may be central to how Hsp70 acts to triage the fate of bound-clients. Prolonged residence in an unfolded, Hsp70-bound state has been shown to promote ubiquitination by the co-chaperone CHIP (carboxyl terminus of Hsc70-interacting protein) and biases clients towards degradation rather than refolding^36–39^. Thus, although active refolding by Hsp70 during heat stress is possible *in vitro*^5,17,32^, our results suggest that refolding may be disfavoured *in vivo* during periods of prolonged stress where minimising chaperone engagement and preventing futile cycles may be advantageous. Together, our findings highlight the remarkable adaptability of the human Hsp70 system to recognise misfolded clients across distinct conformational states and actively remodel their folding landscapes to drive productive refolding to the native state.

## Acknowledgments

We would like to thank the technical officers and staff of Molecular Horizons for their support. Special thanks to Dr Lisanne Spenkelink and Dr Dezerae Cox for their insights and discussions regarding this study. This work was funded by the Australian Research Council (DP220103466). AMvO and HE acknowledge funding from the National Health and Medical Research Council (APP1197069, APP1194872).

## Author contributions

**BS**: Conceptualization (equal); Methodology (lead); Investigation (lead); Data curation (lead); Formal analysis (lead); Visualization (lead); Writing – original draft (lead); Writing – review and editing (equal). **NKA**: Investigation (supporting); Conceptualization (supporting); Writing – review and editing (equal). **SM**: Investigation (supporting); Writing – review and editing (equal). **NRM**: Investigation (supporting); Formal analysis (supporting); Conceptualization (supporting); Writing – review and editing (equal). **AMvO**: Conceptualization (equal); Writing – review and editing (equal). Funding acquisition (equal). **HE**: Conceptualization (equal); Writing – review and editing (equal). Funding acquisition (equal).

## Declaration of interests

The authors declare no competing interests.

## Notes

https://doi.org/10.5281/zenodo.20389093

https://doi.org/10.5281/zenodo.20473167

### Competing Interest Statement

The authors have declared no competing interest.

## References

1. Mayer, M. P. Hsp70 chaperone dynamics and molecular mechanism. Trends Biochem. Sci. 38, 507–514 (2013).

2. Mayer, M. P. & Gierasch, L. M. Recent advances in the structural and mechanistic aspects of Hsp70 molecular chaperones. J. Biol. Chem. 294, 2085–2097 (2019).

3. Mayer, M. P. & Bukau, B. Hsp70 chaperones: cellular functions and molecular mechanism. Cell. Mol. Life Sci. CMLS 62, 670–684 (2005).

4. Kellner, R. et al. Single-molecule spectroscopy reveals chaperone-mediated expansion of substrate protein. Proc. Natl. Acad. Sci. U. S. A. 111, 13355–13360 (2014).

5. Imamoglu, R., Balchin, D., Hayer-Hartl, M. & Hartl, F. U. Bacterial Hsp70 resolves misfolded states and accelerates productive folding of a multi-domain protein. Nat. Commun. 11, 365 (2020).

6. Marzano, N. R., Paudel, B. P., van Oijen, A. M. & Ecroyd, H. Real-time single-molecule observation of chaperone-assisted protein folding. Sci. Adv. 8, eadd0922 (2022).

7. Marzano, N. R. et al. Direct single-molecule visualization of Hsp90-mediated relief of a Hsp70-folding block. Preprint at 10.1101/2025.08.25.672245 (2025).

8. De Los Rios, P., Ben-Zvi, A., Slutsky, O., Azem, A. & Goloubinoff, P. Hsp70 chaperones accelerate protein translocation and the unfolding of stable protein aggregates by entropic pulling. Proc. Natl. Acad. Sci. 103, 6166–6171 (2006).

9. Goloubinoff, P. & Rios, P. D. L. The mechanism of Hsp70 chaperones: (entropic) pulling the models together. Trends Biochem. Sci. 32, 372–380 (2007).

10. Rukes, V., Rebeaud, M. E., Perrin, L. W., De Los Rios, P. & Cao, C. Single-molecule evidence of Entropic Pulling by Hsp70 chaperones. Nat. Commun. 15, 8604 (2024).

11. Kampinga, H. H. & Craig, E. A. The HSP70 chaperone machinery: J proteins as drivers of functional specificity. Nat. Rev. Mol. Cell Biol. 11, 579–592 (2010).

12. Zuiderweg, E. R. P., Hightower, L. E. & Gestwicki, J. E. The remarkable multivalency of the Hsp70 chaperones. Cell Stress Chaperones 22, 173–189 (2017).

13. Farruggia, B. & Picó, G. A. Thermodynamic features of the chemical and thermal denaturations of human serum albumin. Int. J. Biol. Macromol. 26, 317–323 (1999).

14. Wang, Q., Christiansen, A., Samiotakis, A., Wittung-Stafshede, P. & Cheung, M. S. Comparison of chemical and thermal protein denaturation by combination of computational and experimental approaches. II. J. Chem. Phys. 135, 175102 (2011).

15. Zhao, L. et al. The Hsp70 Chaperone System Stabilizes a Thermo-sensitive Subproteome in E. coli. Cell Rep. 28, 1335–1345.e6 (2019).

16. Auld, N. K., McMahon, S., Marzano, N. R., Van Oijen, A. M. & Ecroyd, H. Competing chaperone pathways in α-synuclein disaggregation and aggregation dynamics. Protein Sci. 34, e70296 (2025).

17. Sharma, S. K., De Los Rios, P., Christen, P., Lustig, A. & Goloubinoff, P. The kinetic parameters and energy cost of the Hsp70 chaperone as a polypeptide unfoldase. Nat. Chem. Biol. 6, 914–920 (2010).

18. Kim, Y. et al. Efficient Site-Specific Labeling of Proteins via Cysteines. Bioconjug. Chem. 19, 786–791 (2008).

19. Chandradoss, S. D. et al. Surface Passivation for Single-molecule Protein Studies. J. Vis. Exp. 50549 (2014) doi:10.3791/50549.

20. Hadzic, M. C. A. S., Börner, R., König, S. L. B., Kowerko, D. & Sigel, R. K. O. Reliable State Identification and State Transition Detection in Fluorescence Intensity-Based Single-Molecule Förster Resonance Energy-Transfer Data. J. Phys. Chem. B 122, 6134–6147 (2018).

21. Tiwari, S., Fauvet, B., Assenza, S., De Los Rios, P. & Goloubinoff, P. A novel fluorescent multi-domain protein construct reveals the individual steps of the unfoldase action of Hsp70. Preprint at 10.1101/2022.02.17.480908 (2022).

22. Morán Luengo, T., Kityk, R., Mayer, M. P. & Rüdiger, S. G. D. Hsp90 Breaks the Deadlock of the Hsp70 Chaperone System. Mol. Cell 70, 545–552.e9 (2018).

23. Banecki, B. & Zylicz, M. Real Time Kinetics of the DnaK/DnaJ/GrpE Molecular Chaperone Machine Action. J. Biol. Chem. 271, 6137–6143 (1996).

24. Russell, R., Jordan, R. & McMacken, R. Kinetic Characterization of the ATPase Cycle of the DnaK Molecular Chaperone. Biochemistry 37, 596–607 (1998).

25. Van Durme, J. et al. Accurate Prediction of DnaK-Peptide Binding via Homology Modelling and Experimental Data. PLoS Comput. Biol. 5, e1000475 (2009).

26. Ben-Zvi, A., De Los Rios, P., Dietler, G. & Goloubinoff, P. Active Solubilization and Refolding of Stable Protein Aggregates By Cooperative Unfolding Action of Individual Hsp70 Chaperones. J. Biol. Chem. 279, 37298–37303 (2004).

27. Sekhar, A., Rosenzweig, R., Bouvignies, G. & Kay, L. E. Mapping the conformation of a client protein through the Hsp70 functional cycle. Proc. Natl. Acad. Sci. 112, 10395–10400 (2015).

28. Lu, J. et al. Energy landscape remodeling mechanism of Hsp70-chaperone-accelerated protein folding. Biophys. J. 120, 1971–1983 (2021).

29. O’Connor, M. S. et al. Role of Hsp70 chaperone in client-protein folding elucidated by Markov state modeling and NMR restraint-assisted molecular dynamics simulations. Biophys. J. S000634952500534X (2025) doi:10.1016/j.bpj.2025.08.022.

30. Rudiger, S. Substrate specificity of the DnaK chaperone determined by screening cellulose-bound peptide libraries. EMBO J. 16, 1501–1507 (1997).

31. Finka, A., Sharma, S. K. & Goloubinoff, P. Multi-layered molecular mechanisms of polypeptide holding, unfolding and disaggregation by HSP70/HSP110 chaperones. Front. Mol. Biosci. 2, (2015).

32. Goloubinoff, P., Sassi, A. S., Fauvet, B., Barducci, A. & De Los Rios, P. Chaperones convert the energy from ATP into the nonequilibrium stabilization of native proteins. Nat. Chem. Biol. 14, 388–395 (2018).

33. Sharma, S. K., De Los Rios, P. & Goloubinoff, P. Probing the different chaperone activities of the bacterial HSP70-HSP40 system using a thermolabile luciferase substrate. Proteins Struct. Funct. Bioinforma. 79, 1991–1998 (2011).

34. Mashaghi, A. et al. Alternative modes of client binding enable functional plasticity of Hsp70. Nature 539, 448–451 (2016).

35. Lashkari, V. D. & Marszalek, P. E. Denaturation of firefly luciferase at heat shock temperatures captured in silico. Biophys. J. 124, 2442–2452 (2025).

36. Connell, P. et al. The co-chaperone CHIP regulates protein triage decisions mediated by heat-shock proteins. Nat. Cell Biol. 3, 93–96 (2001).

37. Qian, S.-B., McDonough, H., Boellmann, F., Cyr, D. M. & Patterson, C. CHIP-mediated stress recovery by sequential ubiquitination of substrates and Hsp70. Nature 440, 551–555 (2006).

38. Stankiewicz, M., Nikolay, R., Rybin, V. & Mayer, M. P. CHIP participates in protein triage decisions by preferentially ubiquitinating Hsp70-bound substrates. FEBS J. 277, 3353–3367 (2010).

39. Gersing, S. K. et al. Mapping the degradation pathway of a disease-linked aspartoacylase variant. PLoS Genet. 17, e1009539 (2021).

