## Supplementary Information for "Single-molecule visualisation of human Hsp70-driven conformational remodelling during stress"

### Supplementary Figures

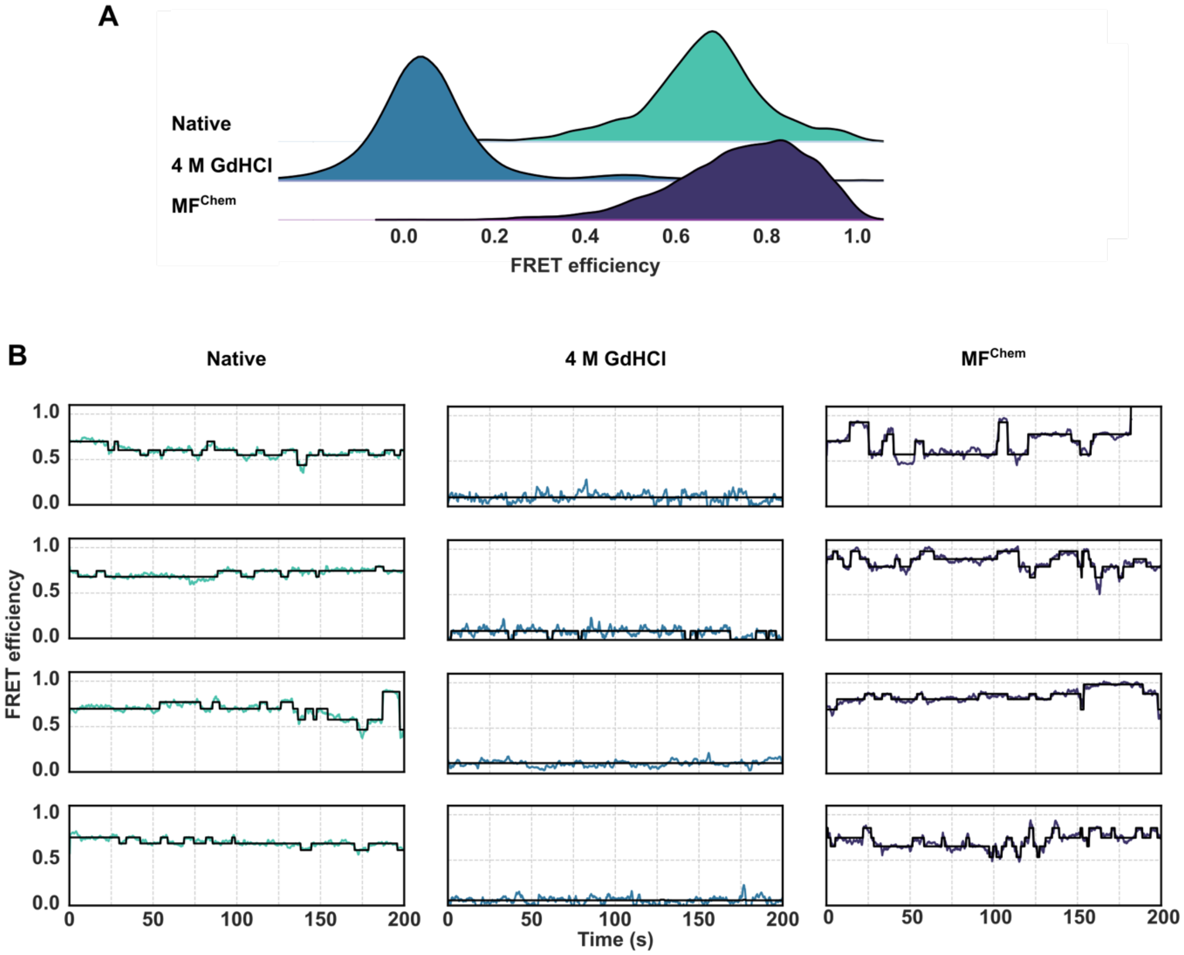


**Figure S1: Incubation with chemical denaturant results in conformational expansion of Fluc$^{IDS}$. (A)** Natively folded Fluc$^{IDS}$ was incubated with 4 M GdHCl. Removal of the denaturant results in Fluc$^{IDS}$ forming a compact-misfolded structure (MF$^{Chem}$). **(B)** Example FRET efficiency traces for Fluc$^{IDS}$ images natively (left), incubated with 4 M GdHCl (centre) or following removal from denaturant (MF$^{Chem}$; right). The idealised trace generated by the Hidden Markov Model fit is indicated in black, whilst the raw FRET efficiency data are depicted in colour.

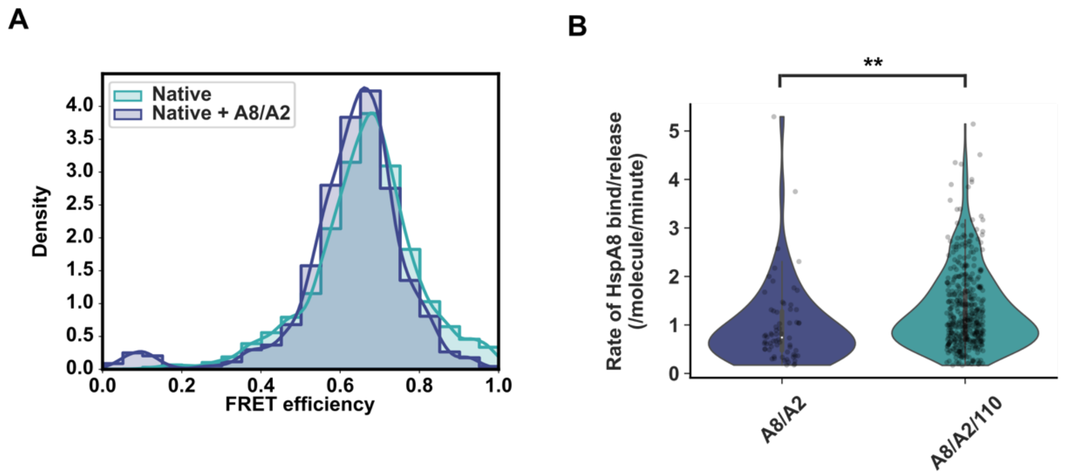


**Figure S2: HspA8 does not bind to native Fluc$^{IDS}$ and folds clients via repeated folding and unfolding events in combination with Hsp110. (A)** Histogram depicting the FRET efficiency of native Fluc$^{IDS}$ incubated alone (Native) or in the presence of 3 μM HspA8, 2 μM DnaJA2 and 5 mM ATP. **(B)** Violin plot depicting the rate of transitions above or below 0.3 FRET, indicative of a HspA8 bind or release event in the presence of HspA8 (3 μM) and DnaJA2 (2 μM) alone (A8/A2), or when supplemented with the complete Hsp70 system (0.5 μM Hsp110; A8/A2/110). An unpaired t-test was performed to determine statistically significant rates of chaperone binding events, with ** denoting significance ($P < 0.01$). Data depict a minimum of 66 bind/release events per treatment. Data from all panels were derived from at least 201 molecules per condition.

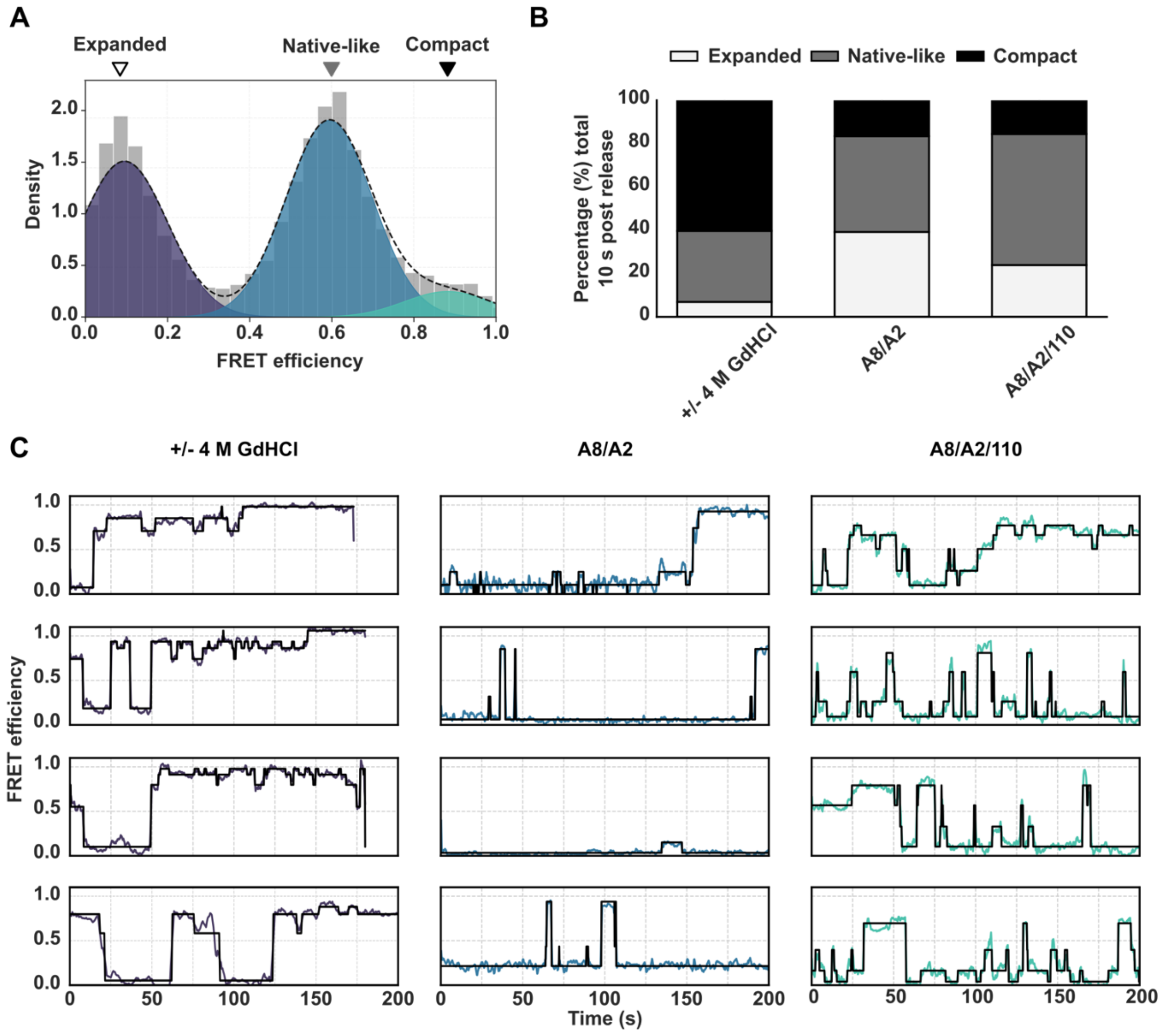


**Figure S3: Gaussian fitting of FRET efficiency data to determine occupancy in different conformational states. (A)** Example FRET efficiency data fit with the sum of 3 Gaussians, with initial estimates for means provided at 0.1, 0.6 and 0.8 to correspond to conformationally-expanded, native-like and compact-misfolded populations, respectively. The area of each Gaussian as a proportion of sum of areas was taken to represent the proportion of molecules occupying the respective state. **(B)** Proportion of FRET efficiency density occupying the three aforementioned states for folding events of $Fluc^{IDS}$ intermittently supplemented with and diluted from 4 M GdHCl (left) or in the presence of A8/A2 alone (centre) or supplemented with Hsp110 (A8/A2/110; right). **(C)** Example traces for $Fluc^{IDS}$ in the presence of or following dilution from 4 M GdHCl (left), in the presence of A8/A2 (centre) or A8/A2/110 (right).

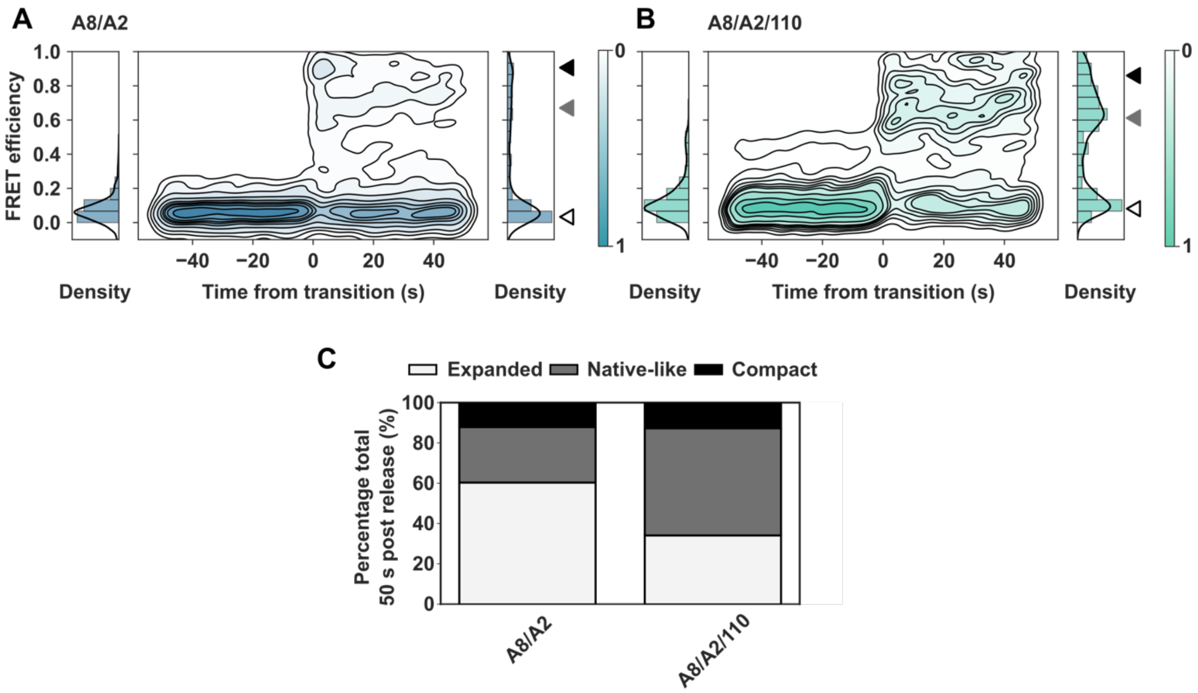


**Figure S4: The presence of the complete Hsp70 system promotes efficient refolding of Fluc$^{IDS}$ to the native state. (A-B)** 2D heatmaps for folding events up to 50 s post release for refolding of MF$^{Chem}$ in the presence of A8/A2 or A8/A2/110, respectively. **(C)** Proportion of density in conformationally-expanded, native-like or compact misfolded conformations up to 50 s post release as determined by fitting with the sum of 3 Gaussians.

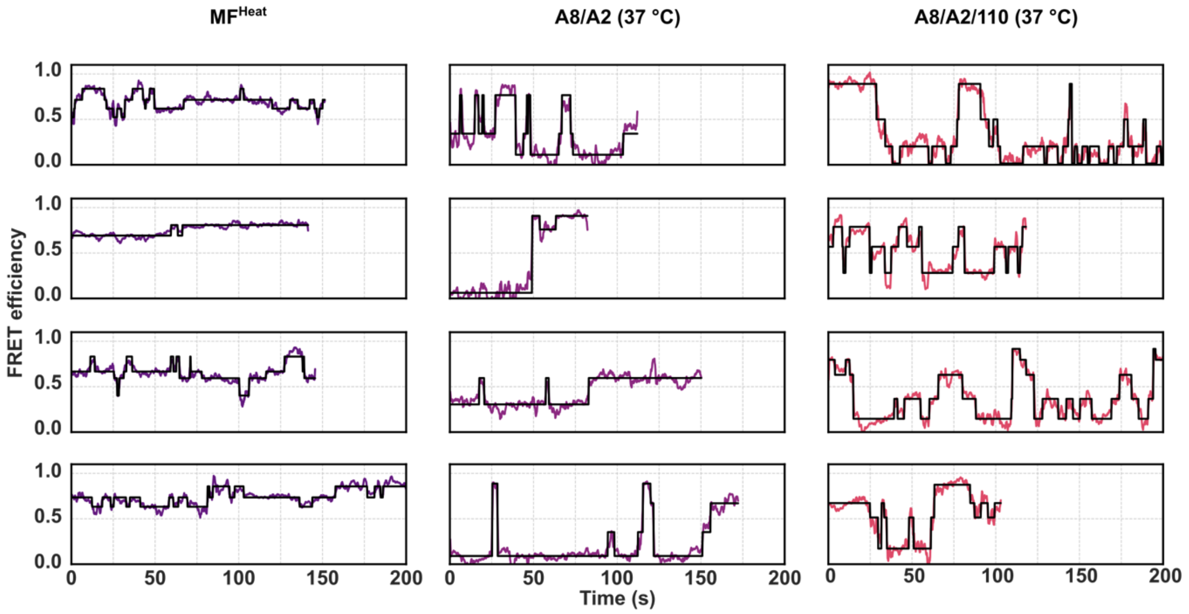


**Figure S5: Example smFRET trajectories of Fluc$^{IDS}$ molecules incubated at 37 °C in the absence (MF$^{Heat}$) or presence of the Hsp70 chaperone system (A8/A2; A8/A2/110).** Raw FRET efficiencies are depicted in colour, with the Hidden Markov Model fit depicted in black.

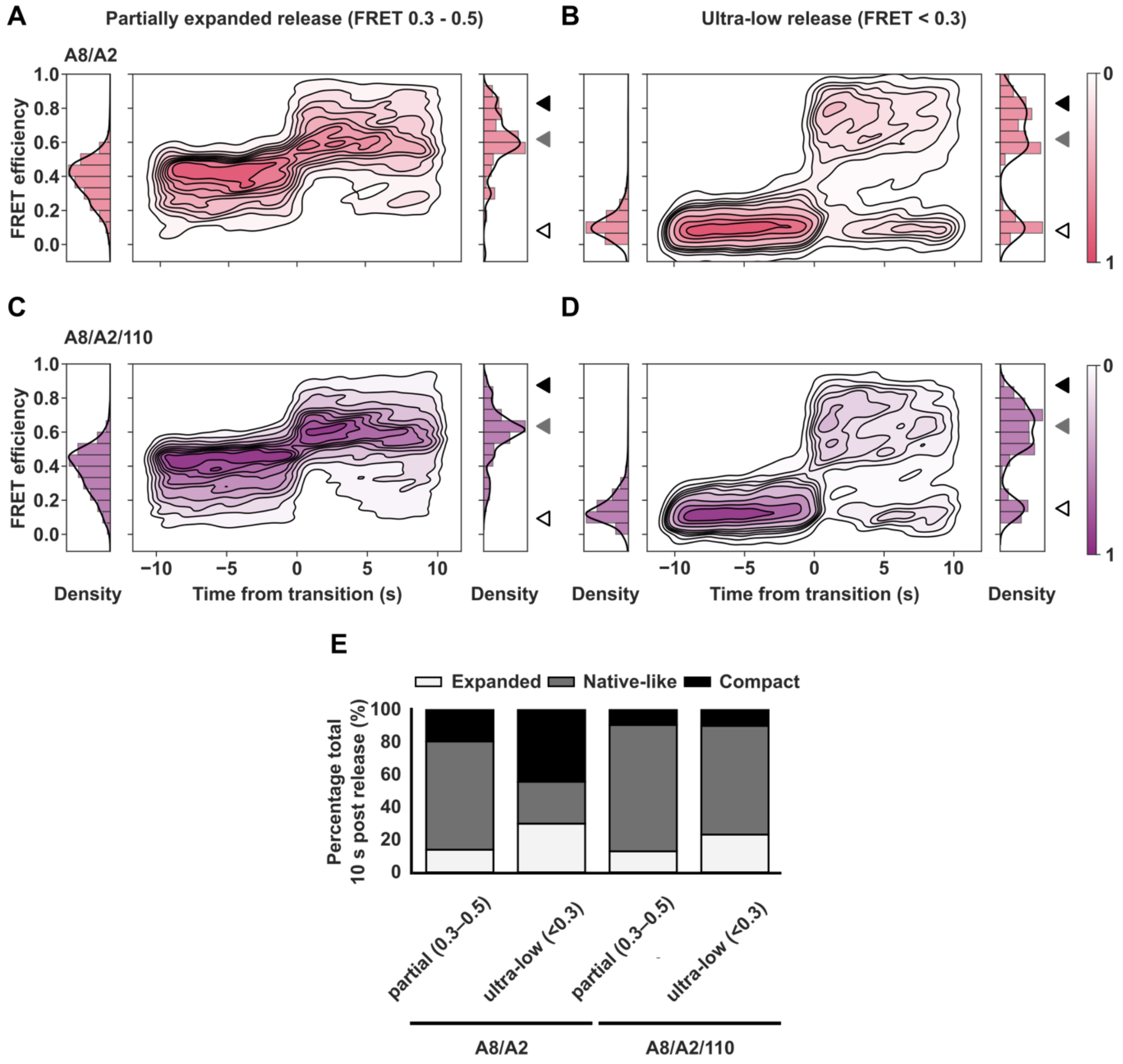


**Figure S6: Folding events are more productive from the partially unfolded Fluc$^{IDS}$ state at 37 °C.** 2D heatmaps depicting synchronised release events from the partially unfolded conformation (FRET efficiency 0.3 – 0.5) **(A, C)** or an ultra-low (FRET efficiency < 0.3) **(B,D)** in the absence and presence of Hsp110. **(E)** Proportion of density found to fit into expanded, native-like and compact misfolded populations as determined by fitting with the sum of 3 semi-constrained Gaussians.

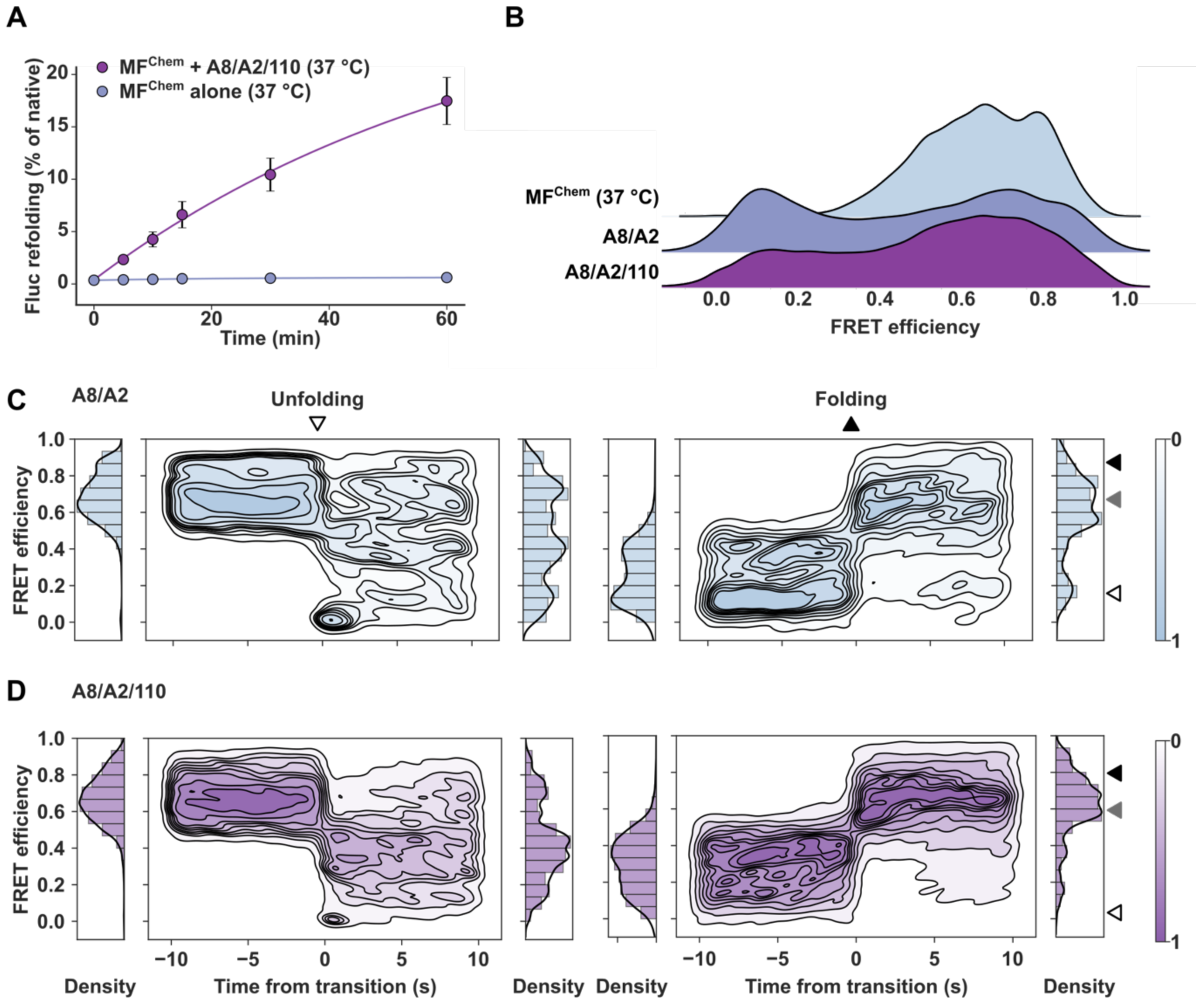


**Figure S7: HspA8 cannot effectively resolve MF$^{Chem}$ at 37 °C. (A)** Enzymatic assay demonstrates limited refolding capacity of HspA8 during heat-stress. Fluc$^{IDS}$ (2 nM) was incubated at 37 °C either in assay buffer alone (MF$^{Chem}$ alone) or in the presence of the complete human Hsp70 system (3 µM HspA8, 2 µM DnaJA2, 0.5 µM Hsp110 with 5 mM ATP; MF$^{Chem}$ + A8/A2/110). Data shown depict mean ± SEM from 3 biological repeats. **(B)** Ridgeline plot depicting smFRET efficiency of Fluc$^{IDS}$ removed from incubation with 4 M GdHCl at 37 °C (MF$^{Chem}$ [37 °C]) and then supplemented with 3 µM HspA8, 2 µM DnaJA2 alone (A8/A2) or with 0.5 µM Hsp110 (A8/A2/110). Data shown are representative of 3 biological repeats with a minimum of 122 molecules per treatment. **(C)** 2D heatmaps depicting unfolding (left) or folding (right) events for MF$^{Chem}$ incubated at 37 °C in the presence of A8/A2 alone or **(D)** supplemented with the complete system (A8/A2/110).

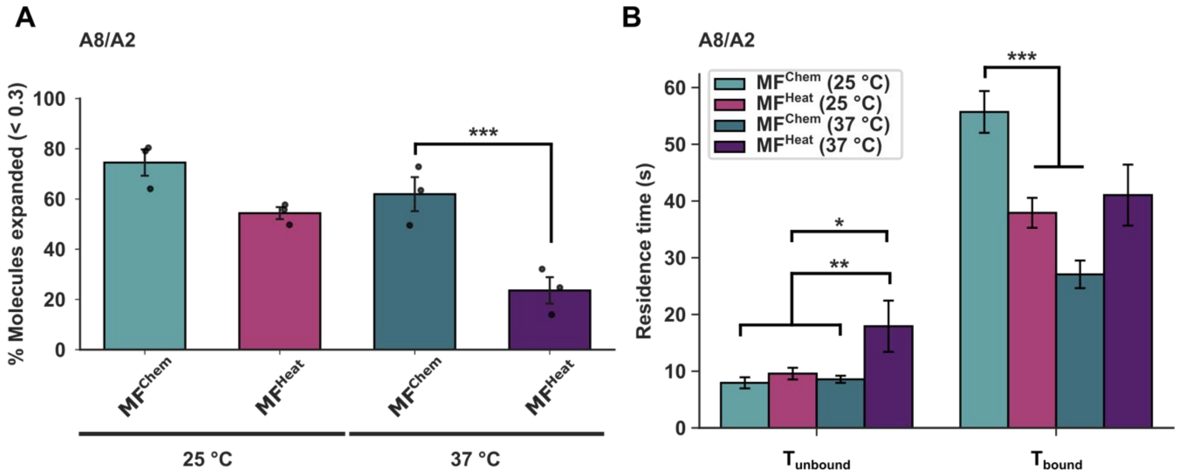


**Figure S8: The nature of the misfolded state dictates the capacity of HspA8 to unfold Fluc$^{IDS}$ into an ultra-low FRET, expanded conformation. (A)** Proportion of Fluc$^{IDS}$ molecules that enter an ultra-low, expanded FRET state (FRET < 0.3) for MF$^{Chem}$ and MF$^{Heat}$ in the presence of A8/A2 at 25 °C or 37 °C, respectively. Data are presented as mean ± SEM from 3 biological repeats. A two-way ANOVA was performed, followed by Tukey's HSD post-hoc test with significance ($P < 0.001$) depicted by ***. **(B)** Mean residence time above ($T_{unbound}$) or below ($T_{bound}$) the ultra-low threshold of FRET 0.3 for MF$^{Chem}$ and MF$^{Heat}$ incubated with A8/A2 for 30 min at either 25 °C or 37 °C. Data are presented as mean ± SEM from 3 biological repeats, with a minimum of 33 events per treatment. A two-way ANOVA followed by Tukey's HSD post-hoc test was performed, where ***, ** and * indicate significance of $P < 0.001$, P < 0.01 and P < 0.05 respectively. Comparisons not indicated were non-significant ($P > 0.05$).

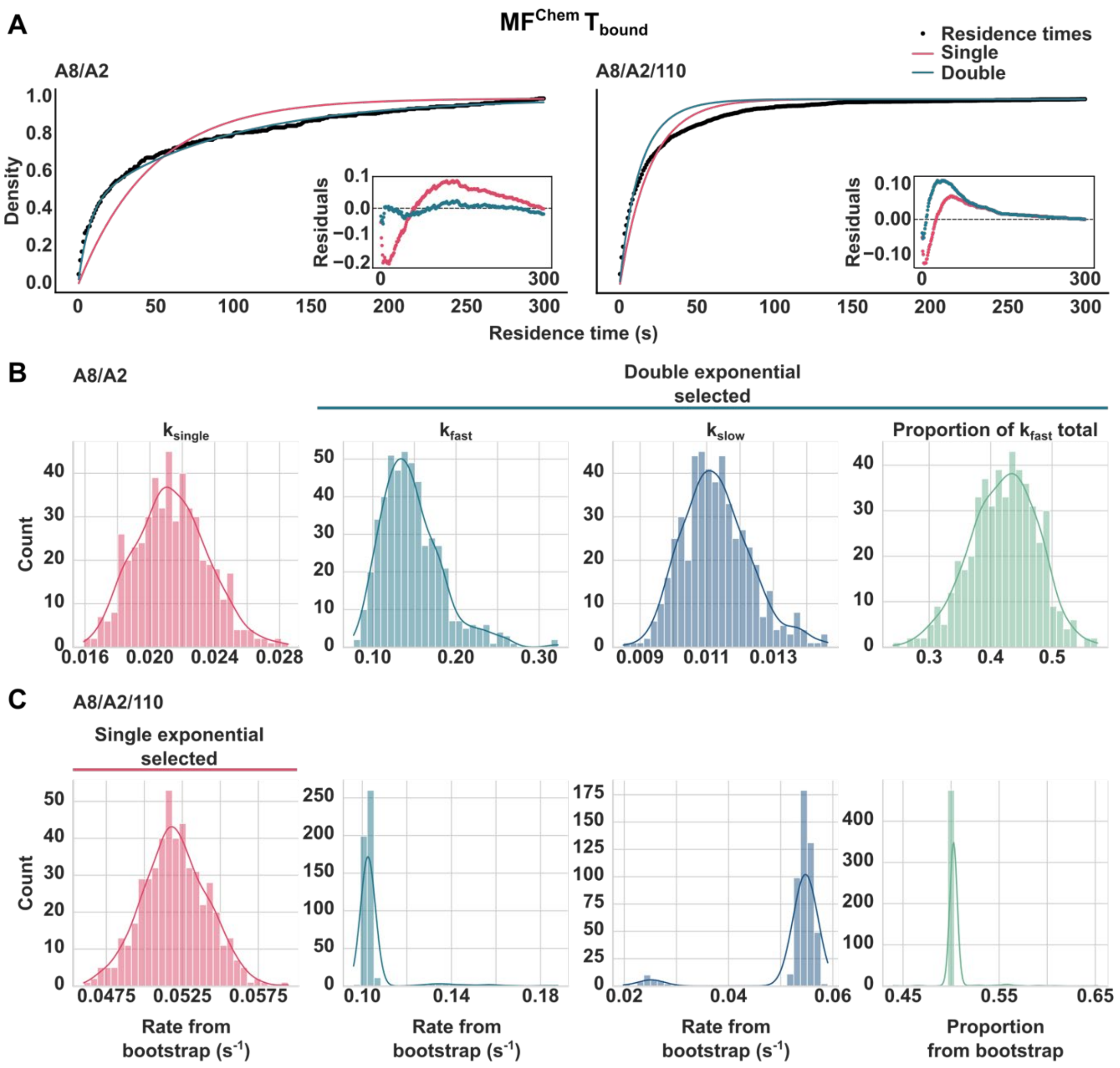


**Figure S9: The stability of the HspA8-MF$^{Chem}$ complex is dictated by two rate constants in the absence of NEF. (A)** The cumulative density function of MF$^{Chem}$ residence times in the $T_{bound}$ state when incubated with A8/A2 alone (left) or supplemented with Hsp110 (A8/A2/110; right). Functions were then fit with either a single (red) or double (blue) exponential function to extract kinetic rates. Residual plots for the single- and double-exponential fits of residence time data depicted in inset figure. Fitting was performed by bootstrap analysis (n = 500), with values for single ($k_{single}$) or double exponential fits ($k_{fast}$, $k_{slow}$ and the proportion of the function dictated by $k_{fast}$ [Proportion $k_{fast}$]) determined for **(B)** A8/A2 or **(C)** A8/A2/110. Final values were obtained from the mean and standard deviation of the bootstrap distributions. Selection of the double-exponential fit was dependent on the combination of decreased residuals and reasonable distributions for proportion of $k_{fast}$ without collapse to a single value as depicted for A8/A2/110. Poor double exponential fits and collapse of the proportion value resulted in selection of the single-rate for these kinetic rates to avoid unnecessary overfitting.

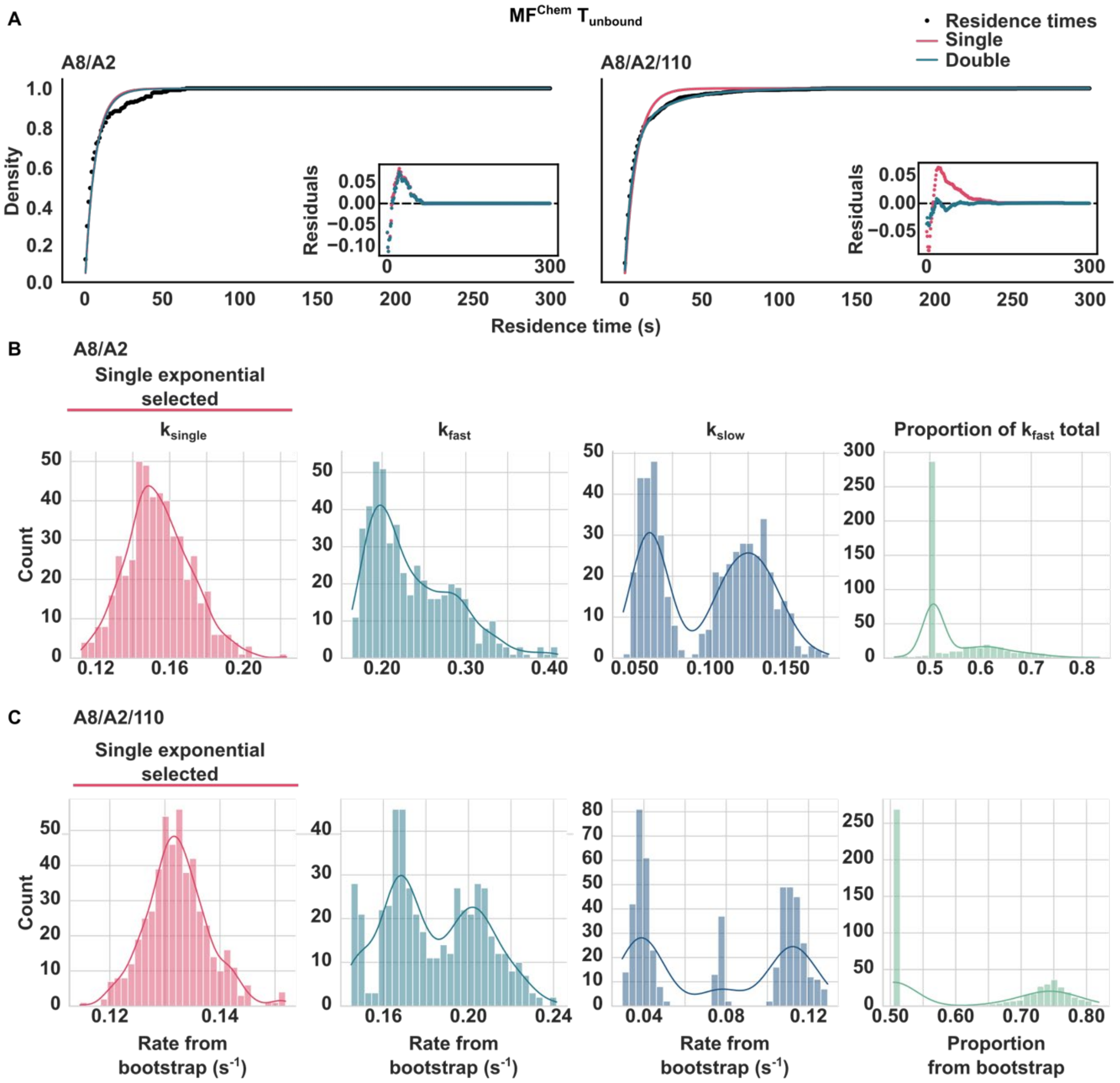


**Figure S10: Kinetics underpinning the unbound state of MF$^{Chem}$ dictated by single rate constant. (A)** Cumulative density functions plots for residence in unbound (*i.e.* FRET > 0.5) conformations for MF$^{Chem}$ in the presence of A8/A2 alone (left) or supplemented with Hsp110 (A8/A2/110; right). Residuals corresponding to single- and double-exponential fits of the data are shown in the inset. Rate constants for the single-exponential ($k_{single}$) and double-exponential ($k_{slow}$, $k_{fast}$) and the proportion of $k_{fast}$ determined for residence in the unbound conformation following 500 bootstrapping iterations are shown for **(B)** A8/A2 alone and for **(C)** the complete chaperone system. Single-exponential fits were selected for both treatments, despite improved residuals, due to poor fitting of the two rate-constants and collapse of the proportion value to 0.5.
